# Chlamydomonas Mutants Null for Chloroplast Triose Phosphate Transporter3 are Metabolically Compromised and Light Sensitive

**DOI:** 10.1101/2022.07.25.501471

**Authors:** Weichao Huang (黄伟超), Anagha Krishnan, Anastasija Plett, Michelle Meagher, Nicole Linka, Yongsheng Wang, Bijie Ren, Justin Findinier, Petra Redekop, Neda Fakhimi, Rick G. Kim, Devin A. Karns, Nanette Boyle, Matthew C. Posewitz, Arthur R. Grossman

## Abstract

Modulation of export of photoassimilates from the chloroplast is essential for controlling the distribution of fixed carbon in the cell and maintaining optimum photosynthetic rates. In this study we identified chloroplast triose phosphate/phosphate translocators 2 and 3 (CreTPT2 and CreTPT3) in the green alga *Chlamydomonas reinhardtii* that exhibited similar substrate specificities but were differentially expressed over the diel cycle. We focused mostly on analyzing CreTPT3 because of its high level of expression and the severe phenotype exhibited by *tpt3* relative to the *tpt2* mutants. Null mutants for CreTPT3 had a pleiotropic phenotype that impacted growth, photosynthetic activities, metabolite profiles, carbon partitioning, and organelle-specific accumulation of H_2_O_2_. These analyses demonstrated that CreTPT3 is a dominant conduit on the chloroplast envelope for the transport of photoassimilate. In addition, CreTPT3 can serve as a safety valve that moves excess reductant out of the chloroplast and appears to be essential for preventing the cells from experiencing oxidative stress and accumulating of reactive oxygen species, even under low/moderate light intensities. Finally, our studies indicate subfunctionalization of the CreTPT transporters and suggest that there are differences in managing the export of photoassimilates from the chloroplasts of Chlamydomonas and vascular plants.

## Introduction

Photosynthetic organisms can absorb excess, potentially damaging levels of light energy during mid-day (when photosynthetic electron transport becomes saturated) and can suffer from extreme damage, especially when experiencing rapid fluctuations in light intensities or when subjected to nutrient limiting conditions and other environmental stresses that impair the productive utilization of excitation energy (Chaux et al., 2017; Saroussi et al., 2017). To cope with excess excitation energy, plants and algae have evolved mechanisms to dissipate this energy through nonphotochemical quenching (NPQ) (Niyogi and Truong, 2013). The excess reductant and energy can also be eliminated by photochemical quenching. While the dominant form of photochemical quenching often involves the use of the electrons/reductant to fix inorganic carbon (Ci), which is directed toward growth or stored in the form of starch and lipids (Krishnan et al., 2015; Ge et al., 2014; Huang et al., 2018), redox equivalents can also be trafficked to other outlets where they are not used for anabolic processes. The main alternative photochemical electron outlets involve the reduction of O_2_ to H_2_O in H_2_O-to-H_2_O cycles that include a (*i*) Mehler-type reaction that is noncatalytic and functions on the acceptor side of PSI (Badger et al., 2000), (*ii*) flavodiiron protein reactions (FLVs, NADPH:flavin oxidoreductase) that catalytically reduce O_2_ on the acceptor side of PSI without generating reactive oxygen species (ROS) (Helman et al., 2003; Chaux et al., 2017), (*iii*) the plastid terminal oxidase (PTOX, plastoquinol: oxygen oxidoreductase) reaction that can use electrons from the PQ pool to reduce O_2_ (Houille-Vernes et al., 2011), and (*iv*) the movement of redox equivalents from the chloroplast to the mitochondrion where they can be used to reduce O_2_ through various electron transport activities.

Mechanisms also exist in which fixed carbon and reducing equivalents can be shuttled between the chloroplast and cytoplasm. Studies with *Arabidopsis thaliana* revealed that the malate shuttle, which involves multiple malate dehydrogenases (MDH) and malate/OAA translocators (OMT), functions in the export of reductant from the chloroplast and the management of redox conditions in the chloroplast (Zhao et al., 2020, 2018). In addition to the malate shuttle, the triose phosphate (triose-P)/phosphate (Pi) translocator (TPT) has been proposed to be involved in moving fixed carbon out of the chloroplast, but can also act as a safety valve for eliminating excess reducing power from the chloroplast (Fliege et al., 1978; Flügge et al., 1989; Lee et al., 2017; Stocking and Larson, 1969; Raghavendra and Padmasree, 2003; Johnson and Alric, 2013). The synthesis of triose-Ps during photosynthetic CO_2_ fixation by the Calvin-Benson-Bassham Cycle (CBBC) is supported by the reducing power/energy (NADPH, ATP) derived from photosynthetic electron transport.

The TPTs reside on the inner chloroplast envelope membrane and can transport triose-Ps (glyceraldehyde 3-phosphate (GAP), dihydroxyacetone phosphate (DHAP)) and the three carbon acid 3-phosphoglycerate (3-PGA) in a counter exchange for cytosolic Pi (Fliege et al., 1978; Lee et al., 2017; Flügge et al., 1989). These transporters belong to a family of plastidic phosphate translocators (pPTs) that function as antiport systems involved in exchanging Pi with phosphorylated C3, C5 or C6 compounds (Flügge et al., 2003). Most angiosperms have two *TPT* genes in their genome, except for various monocots and two dicot families, the Amaranthaceae and the Brassicaceae, which have a single *TPT* gene (Bockwoldt et al., 2019). In plants, cytosolic trioses exported from the chloroplast by the TPTs are used for the biosynthesis of sucrose and other metabolites (Riesmeier et al., 1993) and to drive respiratory activity. In addition to TPTs, plants harbor the three other pPT subfamilies (Fischer et al., 1997; Kammerer et al., 1998; Eicks et al., 2002; Lee et al., 2017; Flügge et al., 1989). These include the glucose 6-phosphate (Glc6P) translocator (GPT), which imports Glc6P into plastids in heterotrophic tissue (Kammerer et al., 1998), the xylulose phosphate translocator (XPT), which plays a key role in coordinating the cytosolic and plastidic pentose phosphate pathways (Eicks et al., 2002), and the phosphoenolpyruvate (PEP) translocator (PPT), which imports PEP into C3 plastids; the PEP can be used for the synthesis of fatty acids, as substrate for the shikimate pathway (Streatfield et al., 1999; Prabhakar et al., 2010), and for the export of PEP from plastids in C4 plants (Häusler et al., 2000).

Over the last few decades, the physiological functions of the TPTs have been examined in some detail. Various plants do not exhibit a strong phenotype if the chloroplast TPT is either eliminated or its level is reduced (Häusler et al., 1998; Walters et al., 2004; Schneider et al., 2002; Riesmeier et al., 1993). A reduction in TPT activity can be compensated for by diverting assimilated carbon into a transitory starch pool that is subjected to accelerated turnover in the light and/or dark (Häusler et al., 1998; Walters et al., 2004; Riesmeier et al., 1993), leading to accumulation of starch degradation products that can be exported from the chloroplast and used in other cellular compartments. Interestingly, a deficiency of the TPT in rice, a plant that uses sucrose stored in the leaves as its major transitory form of fixed carbon, led to severe phenotypic consequences; the plants exhibited reduced photosynthetic rates and decreased levels of both starch and soluble sugars relative to wild type (WT) plants (Lee et al., 2014).

Microalgae have high photosynthetic conversion efficiencies, can thrive in fresh to hypersaline waters and can be metabolically versatile. They have attracted considerable interest worldwide because of their ability to synthesize large quantities of lipids (e.g. for biofuels and food products), starch, pigments and other bioproducts, and can serve in the remediation of wastewater (Khan et al., 2018; Bhatt et al., 2022). The transporters used for moving photoassimilate between the chloroplast and other cellular compartments and the mechanisms and regulation of these transporters in microalgae have not been extensively explored. Developing a more informed understanding of central metabolism in microalgae and the movement of metabolites among compartments can enable additional work on the establishment, regulation, and evolution of metabolic networks in algae and the ways in which algae can be tailored for production purposes and for sustained growth under specific environmental conditions.

In this study, we used Chlamydomonas, a well-established model green algal system that has been extensively used to analyze various physiological processes, to dissect the function of chloroplast TPTs. Of the four putative Chlamydomonas pPTs, we discovered that at least two are ‘genuine’ TPTs (CreTPT2 and CreTPT3) based on yeast liposome transport assays. CreTPT2 was previously reported to be a PPT based on phylogenetic analysis (Bockwoldt et al., 2019). These two TPTs exhibited sub-functionalization that is reflected by their expression levels and temporally and environmentally distinct regulatory patterns. Due to the more severe phenotypes exhibited by *tpt3* relative to *tpt2* mutants, we focused our analyses on Chlamydomonas TPT3 (CreTPT3) which, among the four predicted pPTs, is highly expressed in the light and strongly induced by various environmental stresses. Through a series of physiological analysis, we demonstrated that CreTPT3 is a major conduit on the chloroplast envelope for the trafficking of fixed carbon, sustaining central carbon metabolism, dissipating excess energy, enabling high rates of photosynthetic electron transport, preventing intracellular hydrogen peroxide (H_2_O_2_) accumulation, and balancing redox conditions at the subcellular level.

## Results

### The triose-P/Pi translocator (DMT/TPT) family

Candidate genes encoding triose-P/Pi transporters (TPT) of Chlamydomonas were identified by blasting the Arabidopsis TPT protein (AT5G46110.1) against proteins encoded on the Chlamydomonas genome. The TPT family is the largest within the drug/metabolite transporter (DMT) superfamily in eukaryotes and includes triose-P and sugar-phosphate transporters associated with chloroplasts; many of the family members are still not functionally characterized (Jack et al., 2001; Knappe et al., 2003; Weber et al., 2006). We identified 32 genes encoding potential TPTs in the version v6.1 genome of Chlamydomonas, with four members, CreTPT10 (CreTPT1 in v5.6 genome), CreTPT2 (CreTPT2 in v5.6 genome), CreTPT3 and CGL51 (CreTPT25 in v5.6 genome), predicted to have a transit peptide that would localize the protein to the chloroplast (**Supplementary Table 1**). These four candidates were included in a phylogenetic analysis of plant and algal pPTs that showed that CreTPT3 is a putative TPT, CreTPT2 and CreTPT10 are putative PPTs, and CGL51 is a putative GPT or XPT (Bockwoldt et al., 2019).

To examine the potential substrate specificities of the four putative pPTs, their amino acid sequences were aligned with pPTs from Arabidopsis (AtPTs) (**Supplementary Fig. 1A**). The ability of these transporters to use triose-P/DHAP or 3-PGA as substrate is dependent on five highly conserved amino acid residues (H184, K203, Y338, K359, and R360 in AtTPT1) (Lee et al., 2017; Moog et al., 2020). Of the putative pPTs in Chlamydomonas, only CreTPT3 and CGL51 contain all five of these residues (H170, K189, Y322, K345, and R346 in CreTPT3) (**Supplementary Fig. 1A**). In AtTPT1, residue F262 (F248 in Chlamydomonas in the analogous protein, CreTPT3) is thought to inhibit PEP access to the binding site; this residue is replaced by N in AtPPT1 (PPT) (Lee et al., 2017; Moog et al., 2020). CGL51 has an M at position F248, indicating that unlike CreTPT3, CGL51 might have preference for other substrates. Protein sequence similarity and identity analysis show CreTPT3 and CreTPT2 share the highest and the second highest similarity and identity with AtTPT1, (**Supplementary Fig. 1B**) and are 57% and 49% similar to AtTPT1, respectively. Furthermore, *CreTPT2* and *CreTPT3* have significantly higher transcript accumulation in light relative to Cre*TPT10* and *CGL51* (**Supplementary Fig. 1C, D**). Given the high similarity and high expression levels, CreTPT2 and CreTPT3 were chosen for further study.

### CreTPT2 and CreTPT3 transport properties

We defined the subcellular localization and substrate preferences of both CreTPT2 and CreTPT3. As shown in **Fig. 1A**, both CreTPT2 and CreTPT3 fused to VENUS localized to the chloroplast envelope. To evaluate the substrate specificity of these transporters, each of them was expressed in *Saccharomyces cerevisiae* (yeast) and the resulting recombinant protein was biochemically analyzed using a liposome uptake assay. The Cre*TPT2* and Cre*TPT3* genes were fused at their C termini to a sequence encoding a His tag, codon-optimized, and expressed in yeast (**Fig. 1B**), and total cell membranes were isolated and reconstituted into liposomes (Loddenkötter et al., 1993; Linka et al., 2008). Chloroplast phosphate transporters in vascular plants can catalyze a Pi/Pi homo-exchange in vitro. Both CreTPT2 and CreTPT3 reconstituted in liposomes were able to catalyze the signature Pi homo-exchange, whereas in the absence of a counter-exchange substrate, little Pi uptake was detected (**Fig. 1C** and **D**, left). In contrast, very low Pi uptake rates were observed for liposomes reconstituted with membranes from yeast cells lacking CreTPT2 or CreTPT3 (**Supplementary Fig. 2**), indicating that the introduced transporters were responsible for the detected Pi import activity in the yeast liposomes.

**Fig. 1:**
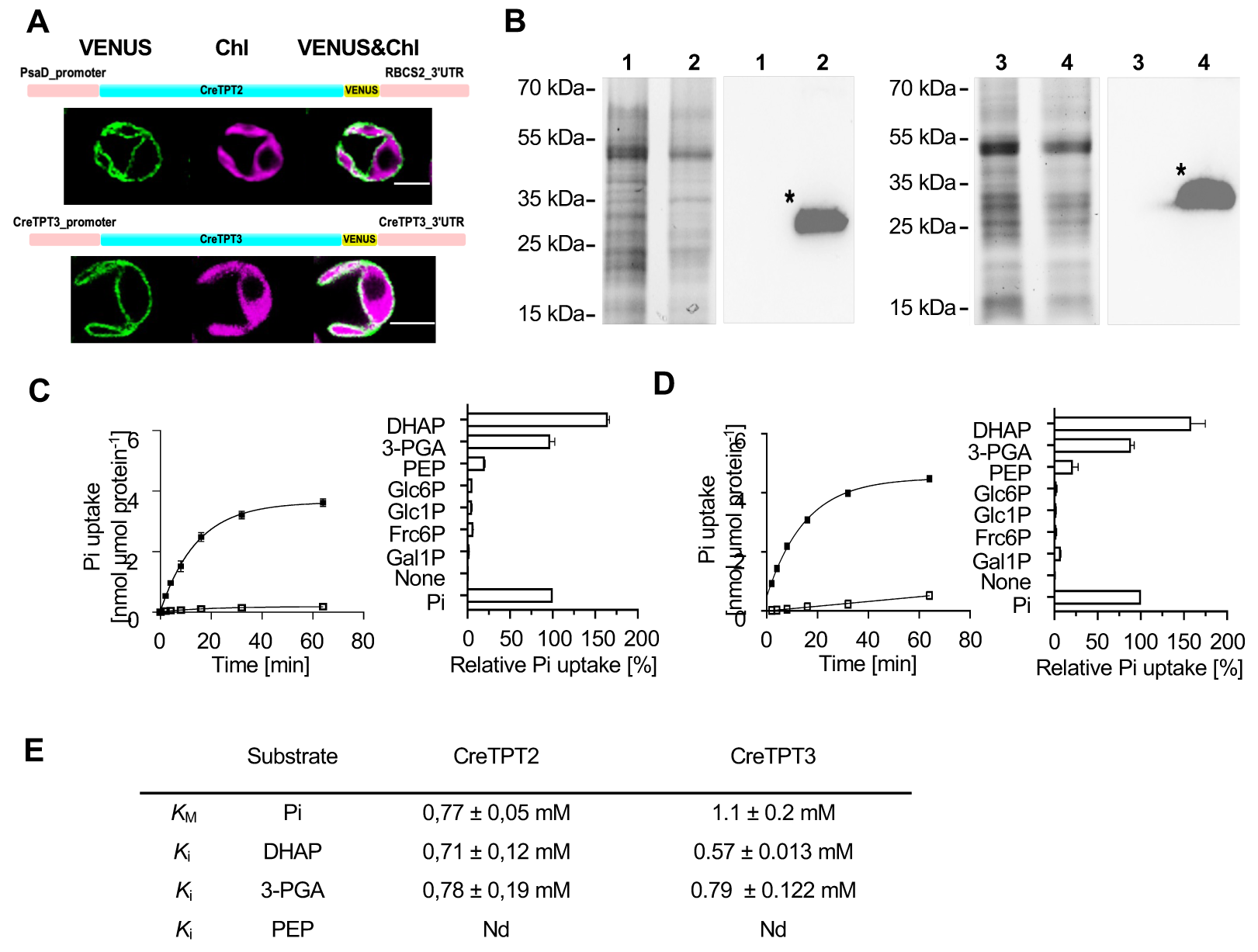
Subcellular localization of CreTPT2 and 3 and in vitro measurements of transport activities. **(A)** Cellular localization of CreTPT2 (upper) and CreTPT3 (lower). The construct for expression of the CreTPT2/3-VENUS fusion protein is shown above the cell images and described in the **Materials and methods**. VENUS fluorescence, green; Chlorophyll (Chl) autofluorescence, magenta. (Scale bar: 5 μm). **(B)** Expression of Cre*TPT2* (1, 2) and Cre*TPT3* (3, 4) in yeast. **Left in each panel**, SDS-PAGE of proteins from total membranes isolated from yeast cells containing the empty vector (1, 3) or expressing His-CreTPT2 (2) or His-CreTPT3 (4). **Right in each panel**, corresponding immunoblot blot detection of recombinant His-CreTPT2 or His-CreTPT3 protein using anti-His antibodies. The calculated molecular masses of the N-terminal His-tagged proteins of CreTPT2 and CreTPT3 were 39, and 37 kDa. (**C**) **Left**, kinetics of Pi exchange by His-CreTPT2 in lipid vesicles. Uptake of Pi (0.25 mM) into liposomes was measured in the presence (◼) or absence (☐) of Pi (30 mM) loaded into the vesicles. **Right**, analyses of substrate specificity of His-CreTPT2. (**D**) **Left**, kinetics of Pi exchange by His-CreTPT3 in lipid vesicles. Uptake of Pi (0.25 mM) into liposomes was measured in the presence (◼) or absence (☐) of Pi (30 mM) loaded into the vesicles. **Right**, analyses of substrate specificity of His-CreTPT3. For the left panels of **C** and **D**, the arithmetic mean (±SD) of three independent experiments (each with three technical replicates) was plotted with respect to the time after initiating the assay. For the right panels of **C** and **D**, liposomes in which His-CreTPT2 and His-CreTPT3, respectively, were incorporated, were preloaded with various substrates (30 mM) and the initial Pi uptake rates (at 0.25 mM external concentration) were determined. Relative velocities were compared to the Pi/Pi homo-exchange, which was set to 100%. The data represents the arithmetic mean (±SD) of three independent experiments (each consisting of three technical replicates). (**E**) Comparison of kinetic constants of His-CreTPT2 and His-CreTPT3. The Michaelis-Menten constant (KM) for Pi uptake was determined using various external Pi concentrations (0.05-5 mM). The competitive inhibition constant (Ki) of Pi uptake (0.25 mM) was measured with increasing competitor concentrations (0.05-5 mM). Liposomes were preloaded with 30 mM Pi as the counter-exchange substrate. The data represent the arithmetic mean +/- SE of three independent experiments. Nd, no competitive inhibitory constant could be measured under the given experimental conditions. DHAP, dihydroxyacetone phosphate; 3-PGA, 3-phosphoglycerate; PEP, phosphoenolpyruvate; Glc6P, glucose 6-P; Glc1P, glucose 1-P; Fru6P, fructose 6-P; Gal1P, galactose 1-P.

To assess the substrate specificity of the CreTPT transporters, the initial rates of Pi uptake into liposomes preloaded with saturating concentrations (30 mM) of various potential counter-exchange substrates were determined. As the right panels of **Fig. 1C** and **D** show, both CreTPT2 and CreTPT3 exhibited the highest activity when DHAP was used as the substrate for the yeast liposome assay. For both transporters, the relative initial velocity for DHAP/Pi exchange was slightly higher than that of 3-PGA/Pi (the relative 3-PGA/Pi exchange was 75% of DHAP/Pi exchange) while Pi uptake into liposomes preloaded with PEP was much lower (**Fig. 1C** and **D**, right panels). Pi import was negligible when CreTPT2 or CreTPT3 liposomes were preloaded with Glc-6-P, Glc-1-P, Fru-6-P, and Gal-1-P (**Fig. 1C** and **D**, right panels). These results show that CreTPT2 and CreTPT3 have almost the same substrate preferences, with both specifically catalyzing the transport of triose-P and 3-PGA across the membrane in exchange for Pi.

Characterizations of the KM and Ki for CreTPT2 and CreTPT3 were performed to determine the affinity of these transporters for the various substrates. CreTPT3 has an apparent Michaelis-Menten constant (KM) of 1.1 +/- 0.2 mM for Pi (**Fig. 1E**), which is comparable to the value obtained for the vascular plant TPT ortholog (Fliege et al., 1978). CreTPT2 has a slightly lower KM for Pi than CreTPT3 (0.77 +/- 0.05 mM) (**Fig. 1E**). However, while the 3-PGA Ki values were comparable for the two transporters, DHAP was more effective in inhibiting the CreTPT3-dependent Pi exchange than CreTPT2-dependent Pi exchange (**Fig. 1E**). These results suggest that CreTPT3 may have a greater specificity for the transport of DHAP than CreTPT2 and that it may be more effective in transporting C3 phosphorylated compounds than CreTPT2. In the case of PEP, no inhibition of the Pi/Pi homo-exchange was observed, even at the non-physiologically high concentration of 5 mM. Overall, the results of these in vitro assays indicate that both CreTPT2 and CreTPT3 have a typical plant-TPT substrate spectrum (Fliege et al., 1978) and can transport Pi, triose-P (DHAP) and 3-PGA in a counter-exchange mode. However PEP might not be a physiologically relevant substrate for either CreTPT2 or CreTPT3, as was shown for apoplast TPT homologues in other organisms (Lim et al., 2010; Moog et al., 2020). Furthermore, the values generated in these analyses may not precisely reflect transport kinetics in vivo since the two transporters used for these in vitro assays were fused to a His-tag at their N terminus, and the yeast liposomes would have a different lipid composition than the chloroplast inner envelop membrane, where these transporters are normally localized.

### Isolation of *tpt2* and *tpt3* null mutants of Chlamydomonas and their impacts on cell growth

To explore the role of CreTPT2 and CreTPT3 in trafficking carbon and potentially reductant across the chloroplast envelope in vivo, CRISPR knockouts of Cre*TPT2* or Cre*TPT3* were generated. We used the CRISPR-Cas9 editing system to disrupt the Cre*TPT2* or Cre*TPT3* genes while at the same time integrating the hygromycin resistance marker gene (*AphVII*) into the edited site (**Fig. 2A, Supplementary Fig. 3** and **4**). Four independent knockouts of Cre*TPT2* (*t2ko1, t2ko2, t2ko3 and 4*) were obtained, with the marker gene inserted into exon 8 (**Supplementary Fig. 3)**. Three independent knockouts of Cre*TPT3* were obtained, including *t3ko1*, with the marker gene inserted into exon 1, and *t3ko2* and *t3ko3*, with the marker gene inserted into exon 7; no CreTPT3 protein was detected in any of these edited strains (**Fig. 2B**).

**Fig. 2:**
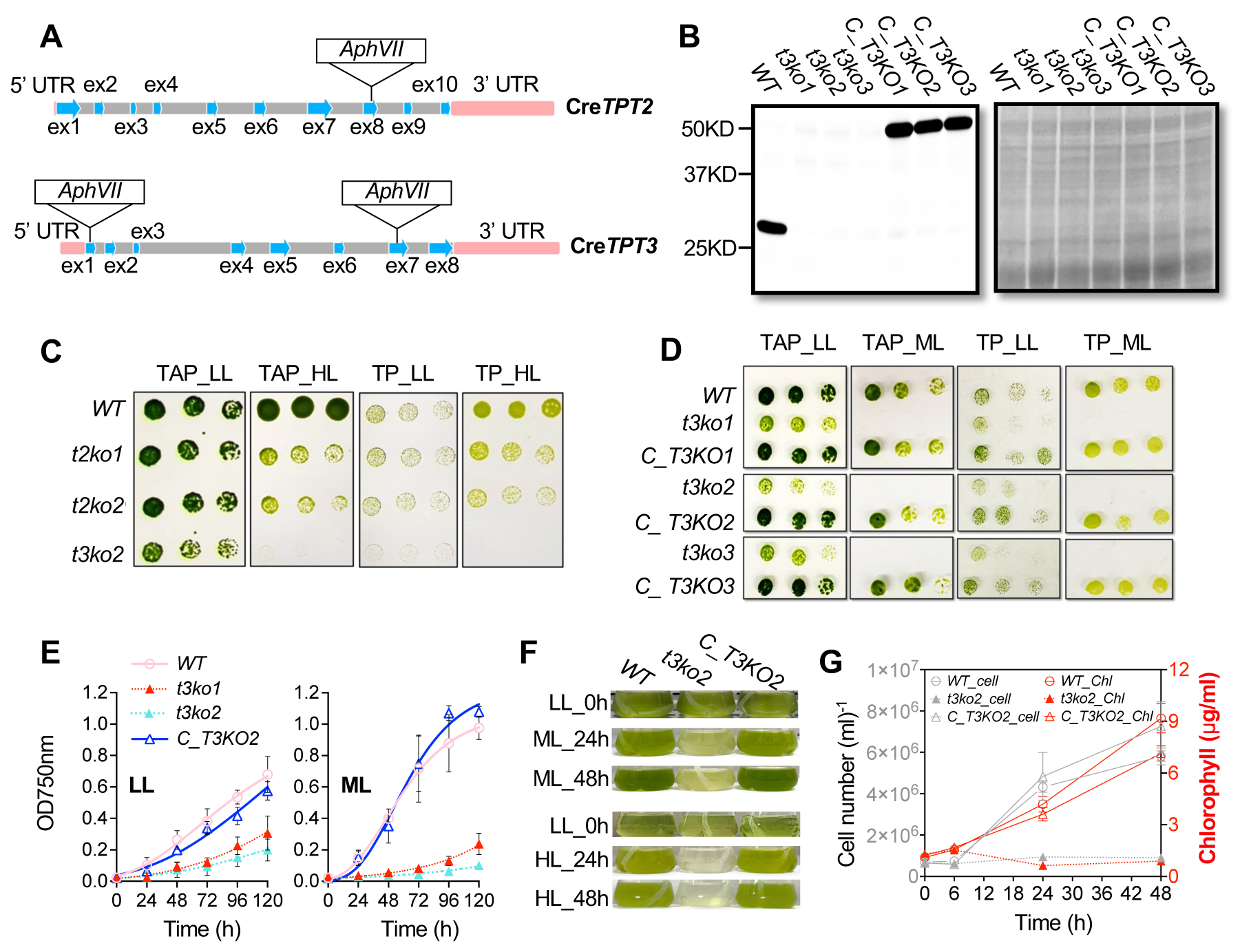
Growth characterization of *tpt* mutants and complemented strains. (**A**) Molecular map of the Cre*TPT2* (upper) and Cre*TPT3* (lower) genes and the positions of the Cas9 targeted sites. Edited site of Cre*TPT2* is located in exon 8, edited sites of Cre*TPT3* are located in exons 1 and 7; *t3ko1* has the marker gene inserted in exon 1 and *t3ko2* and *t3ko3* have the marker gene inserted in exon 7. (**B**) Immunoblot (left panel) of the stained protein profiles (right panel) for wild type (WT), *t3ko1*, *t3ko2*, *t3ko3* and the complemented strains (*C_T3KO1*, *C_T3KO2* and *C_T3KO3*, respectively). (**C**) Growth of WT, *t2ko1*, *t2ko2*, and *t3ko2* at various dilutions (see below) on agar plates incubated under LL and HL for 4 d. (**D**) Growth of WT, *t3ko1*, *t3ko2*, *t3ko3* and the complemented strains (*C_T3KO1*, *C_T3KO2*, and *C_T3KO3*, respectively), at various dilutions (see below) on agar plates incubated under LL and ML for 4 d. (**E**) Growth of the various strains (indicated) in liquid medium under LL (left panel) and ML (right panel). (**F**) Transition of the strains from LL to ML (upper panel) or to HL (lower panel) for the times indicated. (**G**) Cell growth (number) and chlorophyll content of cultures at various times after shifting from LL to ML for up to 48 h. In **C** and **D**, the cells were spotted on agar plates containing TAP or TP medium and maintained under continuous LL, ML or HL conditions. The dilution series used was 1.5, 0.75, 0.375 μg/mL chlorophyll (left to right). For growth in liquid medium, the cells were cultured in TP medium to an initial OD750nm 0.02 in air and under continuous LL before initiating the various growth analyses in LL, ML or HL. Each curve represents the arithmetic mean (±SD) of three independent experiments.

**Fig. 3:**
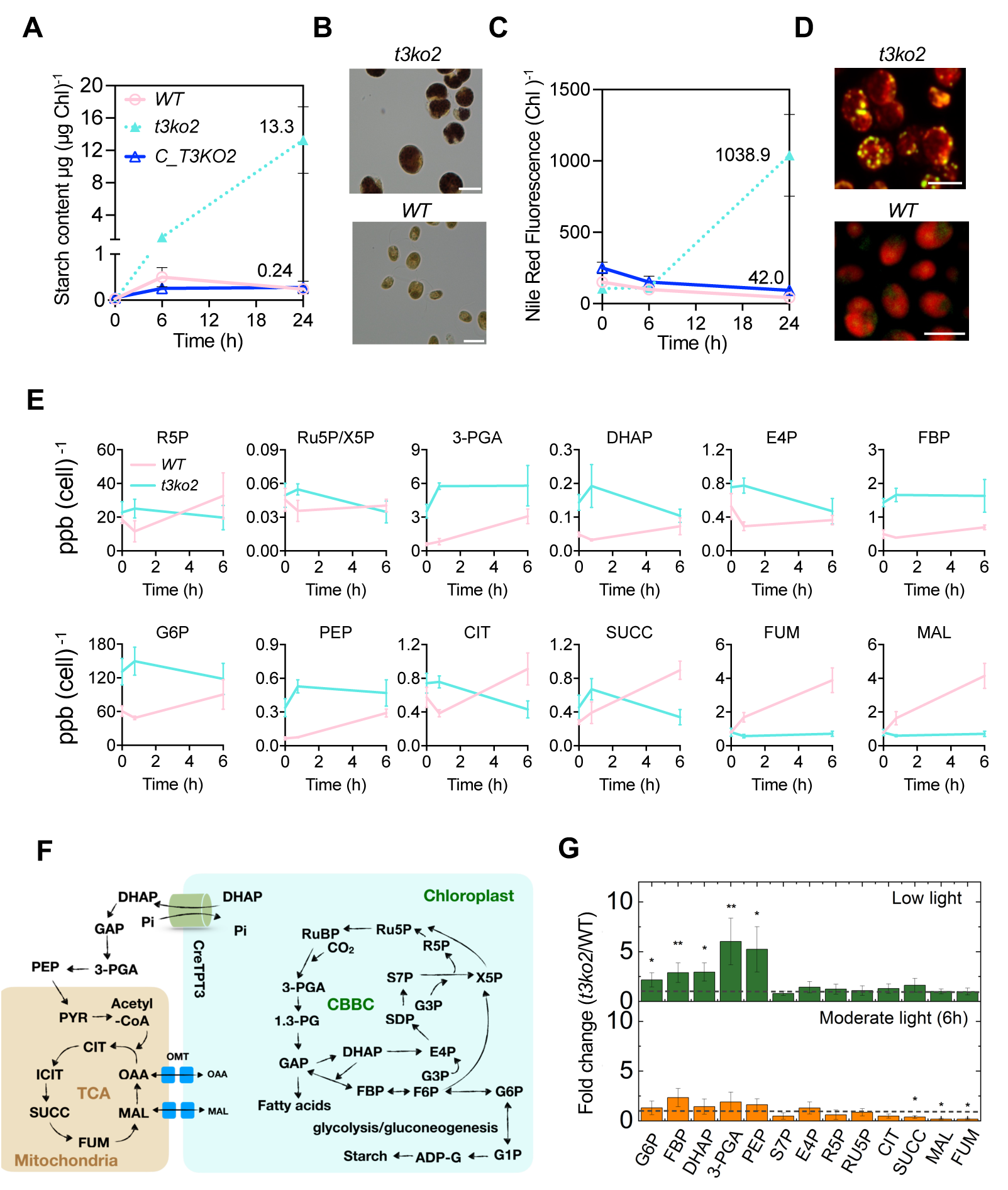
Changes in storage carbon and metabolite levels in WT and *t3ko2* after transitioning cells from growth in LL to ML. (**A**) Starch content in indicated strains following 0, 6, and 24 h of ML exposure. (**B**) Lugol staining of starch in WT and *t3ko2* after 48 h of ML. (**C**) TAG (TAG: triacylglycerol) content in the indicated strains following 0, 6, and 24 h of ML exposure. (**D**) Nile Red staining of TAG in WT and *t3ko2* following 48 h of ML exposure. (**E**) Time course of metabolite accumulation at 0 h (LL), 45 min and 6 h following ML exposure. Data was normalized to cell numbers. Each data point shows the mean and standard error; the data represents three biological replicates for each metabolite. ppb: parts per billion. (**F**) Select metabolic pathways in Chlamydomonas adapted from (Johnson and Alric, 2013). (**G**) Bar graph representation of fold-change for metabolites shown in (**E**); calculated by dividing the mean of pool size in *t3ko2* by that of WT at the respective light levels. A fold change of 1 (no change) is indicated with a dashed line. An asterisk indicates statistically significant difference in pool sizes in *t3ko2* compared to WT (from **E**) (* P<0.05, ** P<0.01, *** P<0.001). Abbreviations: G6P, glucose-6-P; FBP, fructose bisphosphate; DHAP, dihydroxyacetone phosphate; 3-PGA, 3-phosphoglycerate; PEP, phosphoenolpyruvate; PYR, pyruvate; CIT, citrate; SUCC, succinate; FUM, fumarate; MAL, malate; E4P, Erythrose 4-P; R5P, Ribose 5-P; RU5P/X5P, ribulose 5-P/xylulose-5-P; OMT, 2-oxoglutarate/malate transporter.

**Fig. 4:**
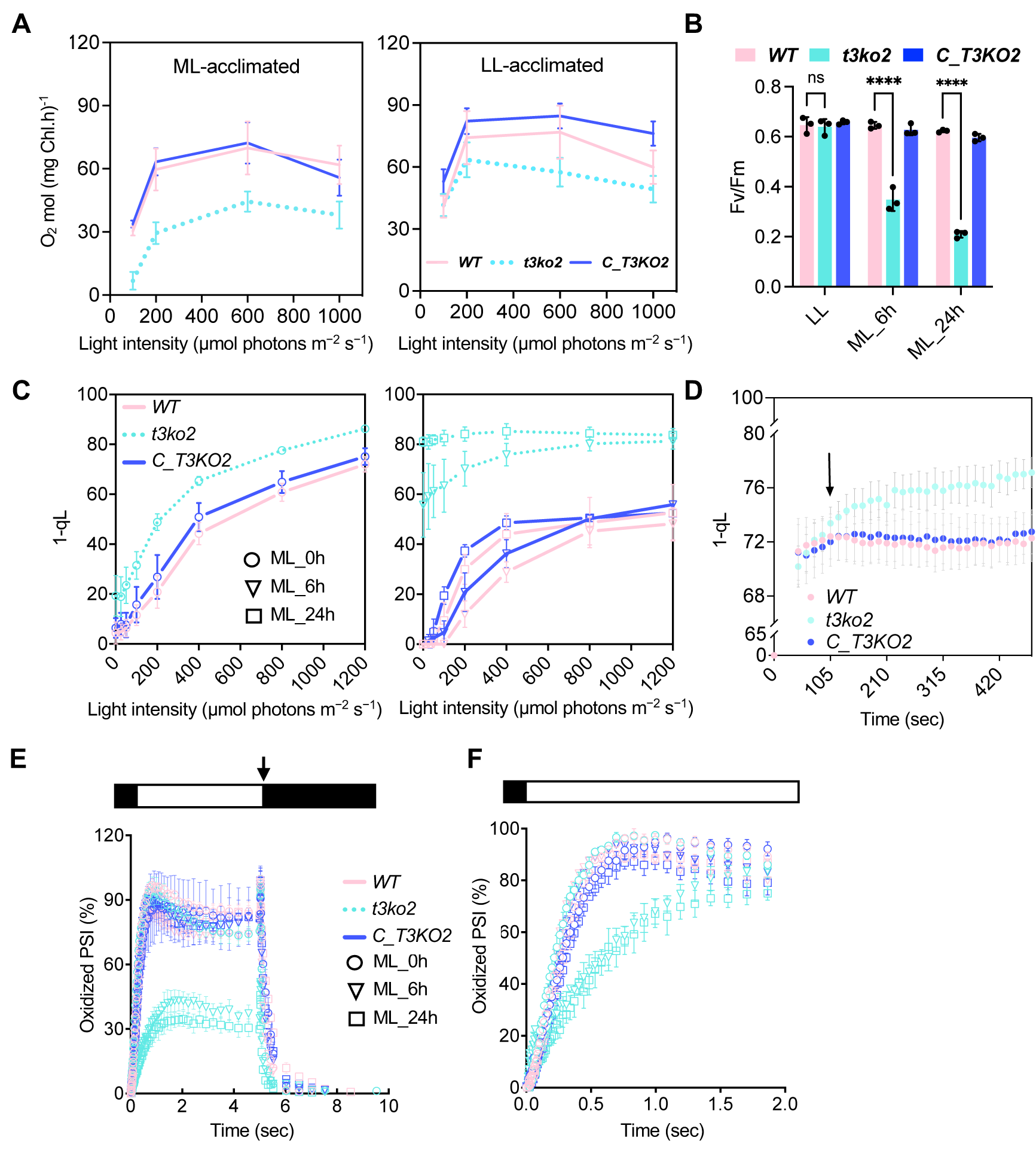
Photosynthetic activities following a transition from LL to ML or to HL. (**A**) Light intensity curve of O2 evolution of LL (left) and ML (right)-acclimated cells for the indicated strains. (**B**) Fv/Fm of WT, *t3ko2* and the rescued strain (*C_T3KO2*) following 0 (LL), 6 and 24 h in ML. (**C**) 1-qL values of LL (left panel) and ML (right panel) acclimated cells exposed to increasing actinic light intensities. (**D**) Kinetics of 1-qL of LL-acclimated cells after illumination at HL (400-450 μmol photons m^-2^ s^-1^) for the times indicated on the x-axis. (**E)** P700 oxidation and reduction kinetics in LL**-** grown cells and after 6 and 24 h in ML. (**F)** Kinetics of P700 oxidation upon dark-to-light transition. P700 measurements were performed in the presence of DCMU and hydroxylamine, as indicated in the **Materials and methods**. Absorbance differences were monitored at 705 nm during continuous illumination with 150 μmol photons·m^-2^ s^-1^ for 5 sec (white box above), followed by a saturating light pulse at 1,500 μmol photons·m^-2^ s^-1^ (arrow) and a 5 sec dark incubation (black box). For panel **E**, the kinetics was normalized by setting maximum oxidation (after light pulse) of WT to 100% and for panel **F**, by setting maximum oxidation (after light pulse) of individual strain to 100%. The data from all panels represent the arithmetic mean (±SD) of three independent experiments. For panel **B**, the asterisks represent significant differences determined by ANOVA tests., ****P < 0.0001.

To elucidate the physiological roles of CreTPT2 and CreTPT3, we first examined growth of the parental WT strain and the knockout mutants under either low light (LL, 30 μmol photons m^-^_2_ s^-1^), moderate light (ML, 250-300 μmol photons m^-^_2_ s^-1^) or high light (HL, 450 μmol photons m^-^_2_ s^-1^). The design of the experiments in which the cells were transferred from one light condition to another is shown in **Supplementary Fig. 5**. The four *tpt2* mutants (*t2ko1, t2ko2, t2ko3 and 4*) were not impacted by growth in LL, but their growth was significantly impaired relative to WT cells under HL on either photoautotrophic (TP) or heterotrophic (TAP) solid agar medium (**Supplementary Fig. 3D, Fig. 2C**). In contrast, the Cre*TPT3* mutants (*t3ko1, t3ko2* and *t3ko3*) exhibited severe growth impairment even in LL, and completely stopped growing under ML and HL, on either TP or TAP agar medium (**Fig. 2C, D**). Due to the striking growth phenotypes caused by the loss of CreTPT3, we focused on investigating the physiological functions of TPT3 by analyzing *tpt3* mutants in more detail in this study. The differences in transcript changes between Cre*TPT2* and Cre*TPT3* under various environmental conditions were examined and are described in the ‘Discussion’ section.

Growth curves were also determined for WT and *tpt3* mutant cells in liquid medium (TP) in LL, ML and HL (**Fig. 2E** and **2F**); the results were in accord with those observed for the solid medium growth assays. Additionally, *tpt3* mutant cells in both LL and ML exhibited an increased cell diameter and formed clusters of cells that appear to be less able to separate following cell division (**Supplementary Fig. 6A**, **B**). To further confirm that the growth phenotypes are a consequence of the *tpt3* knockout, we introduced a wild-type copy of Cre*TPT3* fused to VENUS into the mutant strains (**Fig. 2B**, right). As shown in **Fig. 2D-G**, ectopic expression of the wild-type Cre*TPT3* in the *t3ko2* mutant (*C_T3KO2*) rescued the reduced growth phenotype of the mutant under all light conditions tested in this study (LL/ML/HL).

We also analyzed the chlorophyll content of photoautotrophically grown cells after transferring them from LL to ML and found that *t3ko2* had reduced chlorophyll content relative to WT cells after 24 h in ML (**Fig. 2F-G** and **Supplementary Fig. 7**). Cell numbers and total chlorophyll were quantified following the LL to ML transition. The chlorophyll levels per cell declined in all strains at 24 and 48 h following transfer to ML; however, it was lower by approximately half in *t3ko2* (0.34 µg/10^6^ cells) relative to either WT cells or the *C_T3KO2* rescued strain (both ∼0.72 µg/10^6^ cells) after 24 h of ML (**Supplementary Fig. 7**), with some additional increase in cell density for WT and *C_T3KO2* after 48 h of ML (which might result in some shading). Finally, when mutant cells were transferred to HL for 24 h they became strongly bleached (**Fig. 2F**).

Taken together, these results indicate that the activity of the CreTPT3 transporter is essential for optimal growth over a range of light intensities (LL/ML/HL).

### The *tpt3* mutant exhibits hyper-accumulation of ‘storage’ carbon

To explore the impact of the loss of CreTPT3 activity on carbon partitioning, we quantified carbon storage (starch and lipids) following a transition of WT and the *t3ko2* mutant from LL to ML. Lugol staining showed extensive starch accumulation in the *t3ko2* cells after a 48-h exposure to ML, whereas WT cells were barely stained (**Fig. 3B**). Furthermore, as shown in **Fig. 3A**, there is a ∼55-fold difference in the level of starch that accumulated in *t3ko2* relative to WT cells (13.29 compared to 0.24 µg starch/µg chlorophyll, respectively) after 24 h of illumination in ML. The mutant also accumulated ∼24-fold more lipid than WT and *C_T3KO2* cells (on a chlorophyll basis), as monitored by Nile Red fluorescence, over the same time period (WT and *t3ko2*: 1038 and 41 Nile Red fluorescence/chlorophyll, respectively) (**Fig. 3C, D**). Since the chlorophyll in the mutant on a per cell basis was approximately 50% relative to that of WT cells after 24 h in ML, the accumulated starch and lipid on a per cell basis would be ∼25-fold more starch and ∼12-fold more lipid in the mutant relative to the WT strain. Additionally, mutant cells are much larger and tended to exhibit more aggregation than WT cells (**Supplementary Fig. 6A, B**). These results suggest that the inability to transport triose-P between the chloroplast and the cytosol through CreTPT3 resulted in a repartitioning of photosynthetic assimilates (carbon, reductant, and ATP) toward the synthesis of both starch and neutral lipid.

### Cre*TPT3* deletion leads to accumulation of CBBC/glycolytic/gluconeogenic intermediates

To understand the metabolic consequences of the loss of CreTPT3 on growth under LL and ML, comparative metabolite analyses of WT and the *t3ko2* mutant were performed on cells grown in LL and after shifting them to ML for both 45 min and 6 h. As mentioned above, the level of starch increased dramatically in the mutant relative to WT cells (**Fig. 3A, B** and **Supplementary Fig. 8A**). Pool sizes of various central carbon metabolites, particularly those of the CBBC/glycolysis/gluconeogenesis pathways and intermediates of the TCA/glyoxylate cycle (**Fig. 3F**), were quantified. Data was normalized to both cell number (**Fig. 3E**) and chlorophyll content (**Supplementary Fig. 8B**). The fold-change in the quantity of each metabolite in the mutant relative to WT under LL or at 45 min and 6 h after the switch to ML is given in **Fig. 3G** and **Supplementary Fig. 8C**.

At the time of shifting cells from LL (**Fig. 3E**, time 0 h) to ML, the *t3ko2* mutant had already accumulated a significantly larger pool (2-to 6-fold) of glycolytic/gluconeogenic intermediates [DHAP, 3-PGA, FBP (all three shared with CBBC), G6P, and PEP] compared to WT cells, suggesting that the loss of CreTPT3 resulted in a back-up of these metabolites within the cell; our hypothesis is that these metabolites are accumulating in the chloroplast stroma due to the reduced ability of the mutant chloroplast to export fixed carbon, which is supported by the observed accumulation of starch and TAG in the mutant strain (**Fig. 3A-D** and **Supplementary Fig. 8A**). While the pool sizes of fumarate and malate (metabolites of the TCA/glyoxylate cycle) were similar in both strains under LL, following the transition of the cells to ML, WT cells exhibited a significant increase in the pool sizes of those metabolites while the mutant maintained a lower level, indicating that the loss of CreTPT3 either directly (by supplying precursors) or indirectly (metabolic rewiring) impacts their levels.

Thus, a primary function of CreTPT3 appears to be the export of photosynthetically synthesized reduced carbon, which would drive metabolic processes in the cytoplasm and other cellular compartments, such as the mitochondrion, while enabling the import of Pi into the chloroplast, which sustains ATP synthesis.

### The *tpt3* mutant has reduced photosynthetic activity when grown in ML

To understand the impact of the loss of CreTPT3 activity on PET, we quantified photosynthetic activities following a transition from LL to ML (**Supplementary Fig. 5**). Photosynthetic O_2_ evolution rates (OERs) were measured for WT, *t3ko2* and *C_T3KO2* over a range of light intensities. LL acclimated WT, *t3ko2*, and *C_T3KO2* showed comparable OERs at intensities below 200 μmol photons m^-^_2_ s^-1^, while at saturating light intensities of ≥600 μmol photons m^-^_2_ s^-1^, the OER of *t3ko2* was diminished slightly relative to the WT and the complemented strain (**Fig. 4A**). Upon acclimation of the cells to ML for 24 h, the *t3ko2* mutant consistently displayed lower OERs than WT cells or *C_T3KO2* under all actinic light intensities used (**Fig. 4A**). Additionally, while the Fv/Fm in the mutant grown in LL was comparable to that of WT and *C_T3KO2* (**Fig. 4B**), after exposure of the mutant cells to ML for 6 h, the Fv/Fm declined to about 50% of the WT and *C_T3KO2* levels; the decline in the mutant continued over a period of 24 h in ML, with a 3-fold decrease in Fv/Fm for *t3ko2* compared to that of WT cells (**Fig. 4B**). These results indicate that damage to PSII reaction centers occurs following exposure of the mutant to ML.

To examine the redox state of the photosynthetic apparatus, the pool of electron acceptors downstream of PSII were evaluated. We quantified the photochemical efficiency (qL) of all strains at various light intensities after growth in LL or shifting them to ML for 6 and 24 h (**Fig. 4C**). The photosynthetic parameter qL indicates the redox state of QA, the primary electron acceptor of the PSII reaction center. Assuming that QA and QB are in equilibrium, qL would reflect the redox status of the PQ pool; a lower qL value indicates a more reduced electron transport chain (Kramer et al., 2004). 1-qL positively correlates with the PQ pool redox state. In the LL-acclimated cells, 1-qL was significantly higher in *t3ko2* even at low levels of actinic illumination compared to that of WT cells and the rescued strain. For cells that had been acclimated to ML (6 h and 24 h), 1-qL was 70% of the near maximum value for *t3ko2*, even under relatively low actinic light conditions (e.g. 100 μmol photons m^-^_2_ s^-1^), indicating a highly reduced PQ pool. In contrast, for WT and the complemented strain at the same light intensity, the 1-qL value attained only ∼20% of the maximum value (**Fig. 4C**). These results suggest that the PQ pool is much more reduced in LL-acclimated *t3ko2* relative to WT and *C_T3KO2* (**Fig. 4C**), and that there is a much more pronounced reduction of this pool at all actinic light intensities in ML grown *t3ko2* mutants. These results suggest a limitation in electron flow downstream of the PQ pool in both LL and ML grown *t3ko2*.

Moreover, we analyzed how fast PET becomes restricted following the transfer of LL-grown *t3ko2*, *C_T3KO2* and WT cells to HL (400-450 µmol photons m^-^_2_ s^-1^) by monitoring changes in the redox state of the PQ pool. We observed that the PQ pool in *t3ko2* began to be more reduced than that of WT cells and *C_T3KO2* after 105 s of elevated illumination (**Fig. 4D**), and that this was even more pronounced at longer times in HL. These results suggest that PET can be rapidly limited by diminished triose-P export from the chloroplast; elimination of CreTPT3 has a strong impact on PET activity.

To determine if PSI was also impacted in the mutant cells, we analyzed PSI/P700 oxidation/reduction kinetics following exposure of LL-grown cells to ML in the presence of DCMU (20 µM) and hydroxylamine (1 mM) to block a contribution of electrons from PSII. Levels of photo-oxidizable P700 following exposure to actinic light for 5 sec were similar in all LL-acclimated strains (**Fig. 4E**). However, the level of photo-oxidized P700 in ML-acclimated cells declined in the mutant to ∼40-50% of that in WT and *C_T3KO2* cells (**Fig. 4E**), indicating that PSI is more reduced in the *t3ko2* mutant after ML exposure. As shown in **Fig. 4F**, the oxidation rate of P700 following a dark to light transition was much slower in the ML-acclimated *t3ko2* compared to that of WT. These results suggest that the mutant has a diminished level of available electron acceptors on the acceptor side of PSI (relative to WT and *C_T3KO2*) after growth in ML.

### CreTPT3 inactivation dramatically affected accumulation and distribution of compartmentalized H*_2_*O*_2_*

To investigate the relationship of CreTPT3 activity to oxidative stress, we assayed ROS production in the mutant using the fluorescent probe CM-H2DCFDA, which upon exposure to increasing ROS levels is converted to the green-fluorescent molecule dichlorofluorescein (DCF). DCF fluorescence was visualized by confocal microscopy. As shown in **Fig. 5A** and **B**, ROS levels in the *t3ko2* mutant were markedly increased (∼3-fold) after 48 h in ML, while little difference in ROS levels was detected in WT cells.

**Fig. 5:**
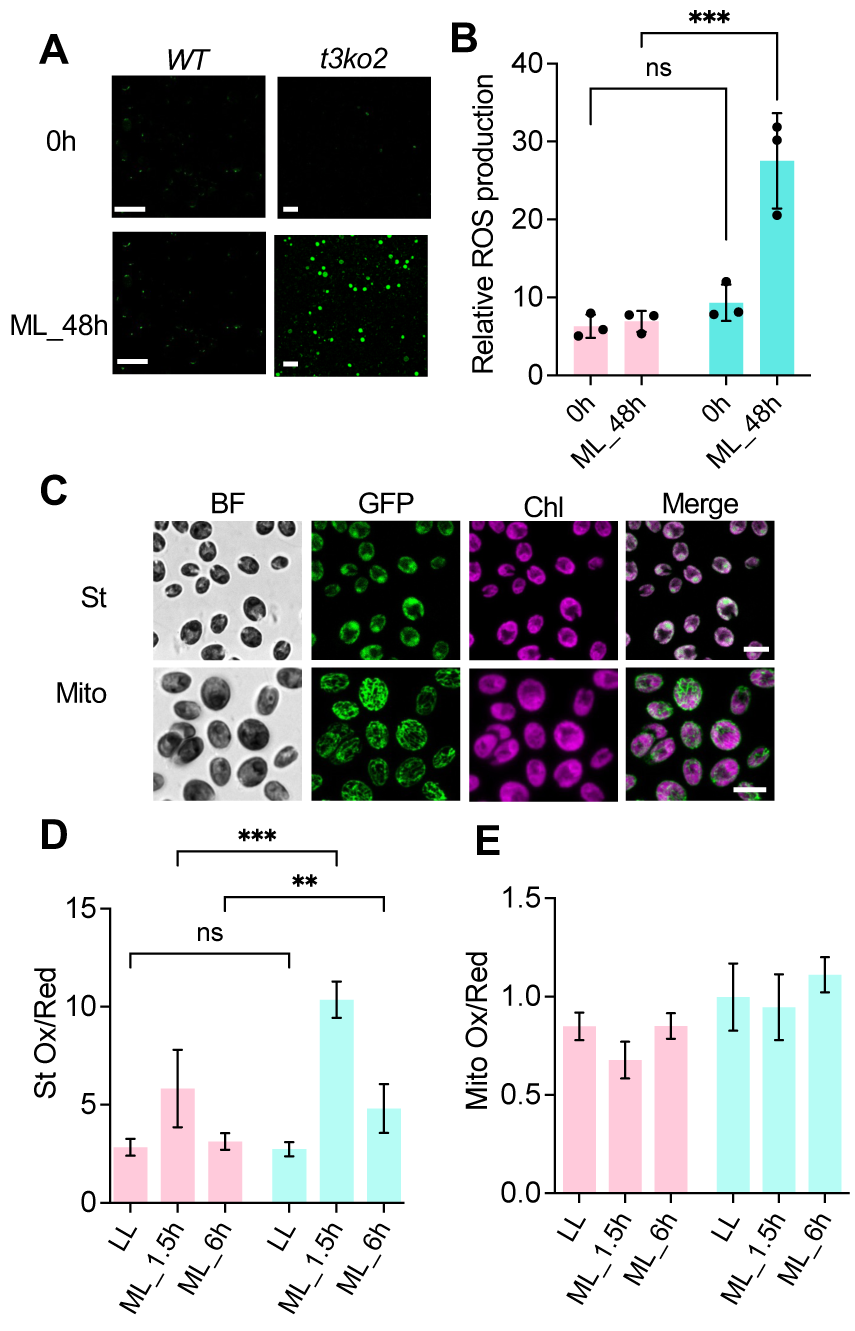
Measurement of intracellular ROS in *t3ko2* and WT upon transition from LL to ML. (**A**) ROS levels were evaluated by CM-H2DCFDA fluorescence of WT and the *t3ko2* mutant in LL and following exposure to ML for 48 h. Scale bar: 20 μm. (**B**) Quantitation of data in **A**. Shown are mean values from three independent experiments, error bars represent standard deviation. roGFP2 protein was targeted to the chloroplast stroma (**C**, upper panel) and mitochondrion (**C**, lower panel). Scale bar: 10 μm; BF: bright field; roGFP2 fluorescence: green; Chlorophyll (Chl) autofluorescence: red. (**D-E**) Monitoring redox levels in stroma (**D**) and mitochondrion (**E**) after exposure of WT and the *t3ko2* mutant for 1.5 and 6.0 h of ML. Shown are mean values from three independent experiments, 3-4 random figures were analyzed for each experiment, error bars represent standard deviation. For panel **B** and **D**, the asterisks represent significant differences determined by ANOVA tests., **P < 0.005, ***P < 0.001.

For an alternative, dynamic method for evaluating redox changes in the chloroplast, we used the redox sensitive green fluorescent protein roGFP2, which was targeted to the chloroplast stroma (**Fig. 5C**, upper panel) and mitochondrion (**Fig. 5C**, lower panel). roGFP2 monitors ratiometric redox changes of glutathione, which reflects cellular ROS levels (Dorion et al., 2021; Vevea et al., 2013). LL-acclimated WT cells and the *t3ko2* mutant exhibited similar levels of chloroplast roGFP2 oxidation (**Fig. 5D**). Upon transfer of these cells to ML, the mutant showed an increase in chloroplast oxidative conditions, with a 5.0-fold increase after 1.5 h, and a 2.0-fold increase after 6 h, which is 1.8-and 1.6-fold higher than the values measured in WT cells (**Fig. 5D**). Additionally, as triose-P is exported to the cytosol by CreTPT3, it could potentially be further metabolized and donate redox equivalents to the mitochondrial electron transport chain and alter mitochondrial ROS production. Therefore, we also measured mitochondrial redox levels at different light intensities in both WT and the *t3ko2* mutant using the roGFP2 sensor targeted to the mitochondrion. Upon a LL to ML shift for 1.5 and 6.0 h, neither WT nor *t3ko2* displayed a significant change in fluorescence from the mitochondrial targeted roGFP (**Fig. 5E**). These results suggest that the mitochondrial redox level is maintained after shifting either LL acclimated WT or *t3ko2* cells to ML. Overall, the inability to export triose-P through CreTPT3 markedly increased the level of oxidative stress in the chloroplast but not in the mitochondrion.

We also determined if the ROS accumulated in *t3ko2* is H_2_O_2_ and whether this molecule shows differential accumulation in the different subcellular compartments; the analysis was based on the use of a hypersensitive sensor of H_2_O_2_, roGFP2-Tsa2ΔCR, which was previously used for studies with Chlamydomonas (Niemeyer et al., 2021). In this analysis, we monitored real-time accumulation of H_2_O_2_ in the stroma, cytosol, mitochondrion, and nucleus (**Fig. 6A**) following a 20 min exposure of the cells to either HL or very low light (**Fig. 6B-E**). The *t3ko2* stromal H_2_O_2_ level increased within 2.5 min of the light exposure and attained a 1.4-fold increase after 20 min of illumination with HL compared to the initial level in LL (**Fig. 6B**). The stromal H_2_O_2_ levels in WT cells showed little change after being shifted to HL, however, the level declined when the cells were shifted to 10 μmol photons m^-^_2_ s^-1^, or very low light. Notably, after 20 min in HL, the stroma of the *t3ko2* mutant accumulated ∼1.6-fold more H_2_O_2_ than that of the WT cells; *t3ko2* mutant cells in very low light accumulated a similar amount of stromal H_2_O_2_ as WT cells after exposure to HL (**Fig. 6B**). The cytosolic probe also responded rapidly, with a 1.2-fold increase for WT and a 1.4- fold increase for the *t3ko2* mutant (**Fig. 6C**); note that the initial levels of H_2_O_2_ prior to the transfer of the cells to HL or very low light were much lower in the WT cells than in the mutant. The *t3ko2* cytosolic H_2_O_2_ level was elevated relative to the level in WT cells by 2.1-fold after 20 min of illumination in HL (**Fig. 6C**). We did not observe a significant change in H_2_O_2_ levels in the mitochondrion for either the WT or the *t3ko2* mutant after the cells were shifted to the higher light intensity (**Fig. 6D**). Finally, the *t3ko2* mutant already accumulated much higher levels of H_2_O_2_ in the nucleus in LL compared to that in the WT cells, although both mutant and WT cells did not show significant changes in nuclear H_2_O_2_ levels after HL exposure (**Fig. 6E**).

**Fig. 6:**
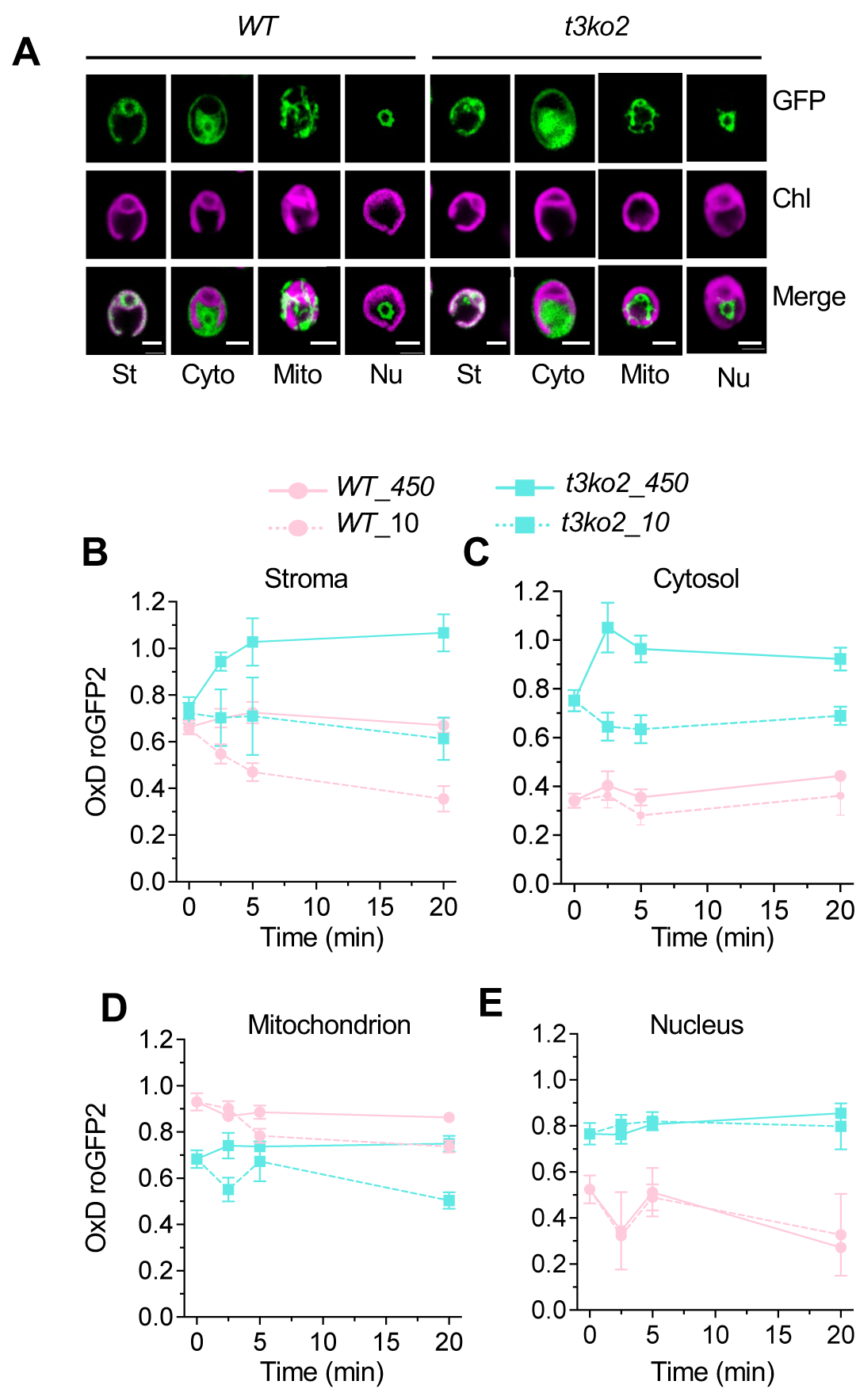
Real-time monitoring of H_2_O_2_ compartmentalized distribution in *t3ko_2_* and WT upon transition from LL to HL. (**A**) The hypersensitive H_2_O_2_ sensor (roGFP2-Tsa2ΔCR) protein was targeted to the chloroplast stroma (St), cytosol (Cyto), mitochondrion (Mito) and nucleus (Nu). Shown are GFP fluorescence, chlorophyll autofluorescence (Chl) and the two signals merged (scale bar: 5 μm). WT and *t3ko_2_* transformant cells, accumulating roGFP_2_- Tsa_2_ΔCR in the stroma (**B**), cytosol (**C**), mitochondrial matrix (**D**), and nucleus (**E**), were acclimated to LL in TP and then were transferred either to HL (450 µmol photons m^−2^ s^−1^, solid line) or to very LL (10 µmol photons m^−2^ s^−1^, dotted line) for _2_0 min. The oxidation state of the sensor was trapped by the addition of NEM and roGFP_2_ fluorescence was measured in a plate reader as previously described (Niemeyer et al., _2_0_2_1). Shown are the mean values from three independent experiments with the error bars representing the standard deviation.

## Discussion

### Cre*TPT2* and Cre*TPT3* genes exhibit different expression patterns

In this study, we discovered that Chlamydomonas contains at least two TPTs that are located on the chloroplast envelope. An earlier report suggested that CreTPT2 was a plastidic PPT (for the transport of PEP) (Bockwoldt et al., 2019), but based on our results, it appears to be functionally more similar to a triose-P transporter. In vitro assays show that CreTPT2 has almost the same substrate specificity as CreTPT3, although it may be less effective in DHAP/Pi exchange (**Fig. 1C-E**). To determine expression patterns of Cre*TPT2* and Cre*TPT3*, the abundance of the Cre*TPT2* and Cre*TPT3* transcripts were analyzed using RT-qPCR and by mining published RNA-seq data over the diurnal cycle, and during nitrogen/sulfur/iron starvation (Zones et al., 2015; Ngan et al., 2015; González-Ballester et al., 2010; Urzica et al., 2013) (**Supplementary Fig. 1C-D,** and **9**). Cre*TPT3* was highly expressed in the light and dark, with significantly higher transcript accumulation than that of Cre*TPT2* and the other genes (Cre*TPT10* and *CGL51*) potentially encoding chloroplast localized pPTs (**Supplementary Fig. 1C-D**). Cre*TPT2* and Cre*TPT3* also responded differentially to abiotic stresses. Cre*TPT3* was strongly induced by nitrogen, sulfur, and iron starvation and upon exposure to HL, whereas the level of the Cre*TPT2* transcript remained almost unchanged under the same conditions (**Supplementary Fig. 9A-D**), suggesting that CreTPT3 plays a more prominent role in exporting triose-P from the chloroplast than CreTPT2, with potentially increasing export from the plastid under HL and nutrient limitation conditions. This hypothesis is supported by the observation that *tpt3* mutants displayed much more severe growth retardation relative to *tpt2* mutants upon exposure to ML or HL (**Fig. 2C**). There is no evidence showing that the *TPTs* from plants are induced by stress/excess absorbed excitation, and expression of the Arabidopsis *TPT* gene appears to even decrease following HL exposure (Weise et al., 2019).

Cre*TPT2* and Cre*TPT3* exhibited distinct expression patterns over the diurnal cycle; the expression of Cre*TPT2* increased rapidly after transitioning from the dark to the light, with peak accumulation after one hour in the light, when the transcript level of Cre*TPT3* was at its lowest (**Supplementary Fig. 1D**). Continued exposure to light led to a decrease in the level of the Cre*TPT2* transcript to near zero while the transcript from Cre*TPT3* steadily increased in the light, reaching a peak in mid-day (**Supplementary Fig. 1D**). The Cre*TPT2* expression pattern suggests that it might play a role in exporting triose-P at the beginning of light period, when the light intensity and photosynthesis are low and low levels of triose-P would be synthesized in the stroma. As the light intensity increases over the course of the day, higher levels of triose-P are synthesized and its trafficking out of the chloroplast for use in other subcellular compartments would likely predominantly involve the activity of CreTPT3. The greater specificity of CreTPT3 than CreTPT2 for transporting DHAP (**Fig. 1E**) may make it more effective than CreTPT2 in transporting C3 phosphorylated compounds. This possibility is congruent with the finding that there is elevated synthesis/accumulation of CreTPT3 mRNA during the day when the light intensity reaches its peak (Zones et al. 2015) and there would be rapid synthesis of the C3 phosphorylated compounds. Overall, the subfunctionalization of the two Chlamydomonas triose-P transporters based on their expression levels, patterns of RNA accumulation over the course of the day and upon nutrient deprivation, and their substrate specificities, may help tune the export of triose-P from the chloroplast with respect to the diurnal cycle and dynamic environmental cues.

### CreTPT3 also potentially serves as a redox valve, transferring reductant to the cytoplasm

It was previously proposed that chloroplast TPTs could catalyze two potential reactions in the light (**Fig. 7**) based on the crystal structure of the red algal TPT (Lee et al., 2017) and in vitro assays using isolated spinach chloroplasts (Stocking and Larson, 1969); both triose-P/Pi, and triose-P/3-PGA exchange across the chloroplast inner envelope membrane. The former reaction can route both carbon skeletons and reductants into the cytoplasm while importing Pi back into the chloroplast for ATP regeneration. The latter reaction would import 3-PGA into the chloroplast in exchange for triose-P (DHAP, GAP), which would serve to transfer reductant from the chloroplast to the cytosol while transferring 3-PGA back into the chloroplast where it can be reduced by the CBBC and stimulate the regeneration of ribulose 1,5-bisphosphate. Indeed, in vitro, CreTPT3 can actively transport both triose-P and 3-PGA in exchange for Pi (**Fig. 1**), indicating that this transporter can serve as both a carbon and ‘reductant shuttle’ which would help sustain photosynthetic electron flow.

**Fig. 7:**
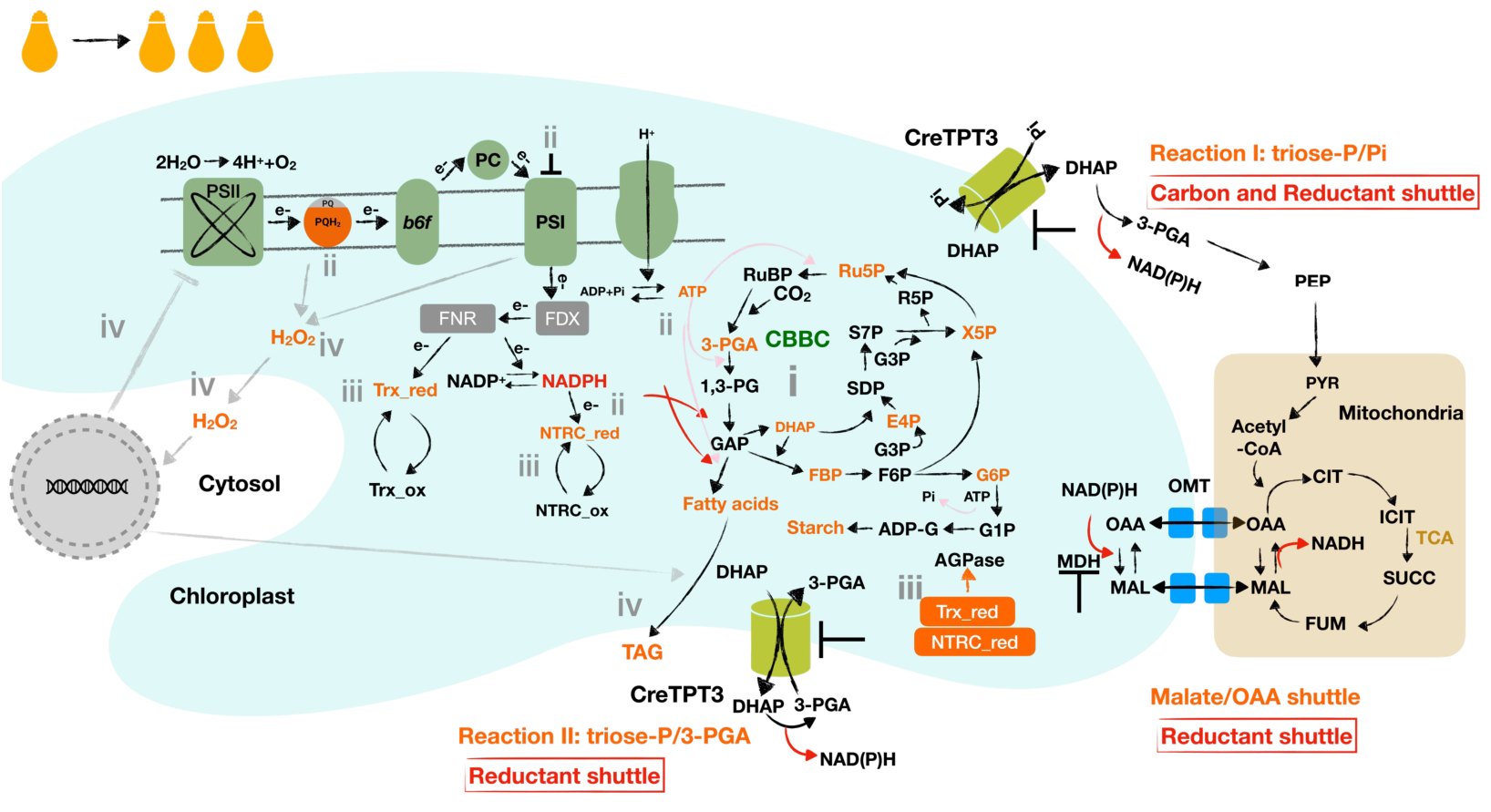
Impact of eliminating Chlamydomonas TPT3 on the chloroplast metabolic landscape. i. reduced triose-P export leads to accumulation of the sugar-P in CBBC and glycolysis pathways (e.g. triose-P, hexose-P, marked in orange), and an elevated ratio of 3-PGA/Pi (which can allosterically activate AGPase activity) would begin to stimulate the synthesis and accumulation of storage carbon; **ii.** CBBC activity diminishes as a consequence of the hyper-accumulation of the precursors with the fixation of CO_2_, leading to elevation of’reductant (NADPH) and energy (ATP) within the chloroplast; the elevated NADPH:NADP^+^ and ATP:ADP ratios, elicit strong feedback on PET causing hyper-reduction of electron carriers (eg: over reduced PQ pool and highly reduced PSI) that slows electron flow across the cytochrome *b6f* complex; **iii.** the highly reduced PET system and the elevated NADPH can actively reduce FDX/TRX and the NTRC systems, respectively; reduced TRX and/or NTRC can activate the AGPase (above the activity elicited by an increase in the 3-PGA/Pi ratio), leading to additional starch accumulation; **iv.** hyper-reduction of PET can also lead to stromal ROS accumulation, that can act as a signal that controls expression of chloroplast and nuclear genes; ROS also cause damage to both PSII and I and leads to neutral lipid accumulation as a consequence of activation of diacylglycerol acyltransferases (DGAT) and phospholipid diacylglycerol acyltransferase (PDAT). Many of these metabolic/acclimatory processes both overlap and are interconnected. Under conditions of extended exposure to ML, the damage in Chlamydomonas *tpt3* mutants can accumulate and lead to cell death; cell death occurs rapidly in HL.

Studies of photosynthetic activities and growth of WT and *tpt3* mutants (e.g. *t3ko2*) in LL, ML and HL support the idea that the mutant is highly compromised in its ability to export fixed carbon and potentially also reductant from the chloroplast. In *t3ko2* exposed to LL (after growth in LL), growth was slow (**Fig. 2E**) and the PQ pool (**Fig. 4C**) was more reduced than in WT cells, while there was little loss of PSII or PSI activities (**Fig. 4A, B, E, F**). These results suggest that there is a reduced rate of PQH2 oxidation. During ML exposure, the mutant stopped growing, and when placed in HL experienced severe bleaching (**Fig. 2F**). The highly reduced PQ pool and PSI reaction center in ML-acclimated *t3ko2* cells (**Fig. 4C, E**, respectively) reflects hyper reduction of PET and the generation of ‘over-flow’ electrons. The phenotypes of the *t3ko2* cells, including an elevated 1-qL (**Fig. 4C-D**), slower oxidation rate of PSI in ML-acclimated cells (**Fig. 4F**), accumulation of storage carbon (**Fig. 3A-D**), an increase in intracellular accumulation of triose-P and 3-PGA (**Fig. 3E**), light dependent damage to the photosynthetic apparatus (**Fig. 4B**) and elevated production/accumulation of ROS (**Fig. 5A, B**), especially in the chloroplast (**Fig. 6B**), indicates a block on the acceptor side of PSI, which reflects the function of CreTPT3 and its central role in fixed carbon export from the chloroplast and for fueling central metabolism. Furthermore, the inability to efficiently transport fixed carbon from the chloroplast in ML and HL would result in diminished CBBC activity, diminished photosynthetic electron transport and the hyper-reduction of the stroma, which would also result in diminished ATP synthesis and NADPH production.

The malate-OAA shuttle represents another route that, under high redox stress, might have partially compensated for the loss of the CreTPT3 by transporting reductant from the chloroplast (schematic in **Fig. 7**). Intriguingly, malate levels in *t3ko2* were 4-fold lower than in WT cells (**Fig. 3E**, **G**). Moreover, expression of the plastid localized malate dehydrogenases (Cre*MDH1* and Cre*MDH5*) was 3-fold to 5-fold lower in the mutant than in WT cells following a transition from LL to ML (**Supplementary Fig. 10D-E**), indicating that the malate-OAA shuttle is likely unable to compensate for a loss of CreTPT3. Inactivation of CreTPT3 appears to have a negative impact on the malate-OAA shuttle, potentially because of the compromised physiological state of the *t3ko2* mutant. Furthermore, a previous study of metabolic flux analysis during heterotrophic growth of Chlamydomonas showed that the CreTPT shuttle(s) is almost 10-fold more active than the malate-OAA shuttle (Boyle et al., 2017).

### TPT deficiency in Chlamydomonas cannot be compensated for by a day/night regime

TPT deficiency in plants can be almost fully compensated for by the starch-mediated night pathways that elicit the breakdown of starch and the export of the breakdown products via the maltose transporter (MEX1) and glucose translocator (GlcT) (Niittylä et al., 2004; Cho et al., 2011). Furthermore, in plants, starch turnover may also be occurring in the light, at the same time as starch is being synthesized (Häusler et al., 1998; Walters et al., 2004). Compared to algae, plant cells appear to display a high plasticity in their capacity to transport fixed carbon between the chloroplast and cytosol. This high degree of plasticity in plants is reflected by the findings: i) most dicots contain a larger number of pPTs (from 5 to 16) (Bockwoldt et al., 2019). For example, Arabidopsis contains 6 pPTs, which includes one TPT, two GPTs, two PPTs and one XPT (Bockwoldt et al., 2019), whereas Chlamydomonas harbors four pPTs, which include two TPTs, one putative PPT (TPT10 in this study) and one putative GPT/XPT (CGL51 in this study); ii) the plant TPTs play an important role in the export of carbon from the chloroplast during the day. However, XPT has been shown to transport triose-Ps and partially compensate for the loss of TPTs under both ML and HL conditions (Eicks et al., 2002; Hilgers et al., 2018b). Hence, it appears that the paths for fixed carbon export in plants are cooperative, with contributions of transporters that use various sugars, sugar phosphates and triose phosphates.

Additionally, the elevated starch content in the *t3ko2* mutant during growth in both LL and ML compared to WT cells suggests that the lesion creates a bottleneck in the export of fixed carbon, which in part becomes stored as starch and lipids (**Fig. 3A-D, Supplementary Fig. 8A**). The conversion of triose-P to starch releases Pi within the chloroplast that can at least partially compensate for the shortage of Pi caused by inactivation of CreTPT3 (Börnke and Sonnewald, 2011). Thus, photosynthetic carbon assimilation can be maintained until the cellular system is completely compromised. In this study, we found that Chlamydomonas *tpt3* mutants exhibited severe growth retardation and the accumulation of starch and lipid in either continuous light (CL) or when experiencing a day/night regime (**Fig. 2D-G** and **8A**). Furthermore, the light-induced electron transport rate (ETR) through PSII in the *t3ko2* mutant maintained on a diurnal cycle was similar to that of WT cells exposed to actinic light intensities of up to 200 μmol photons m^-^_2_ s^-1^, but was approximately 30% lower than that of WT cells at a light intensity of 400 μmol photons m^-^_2_ s^-1^ (**Fig. 8B**). We also observed that diurnally maintained *t3ko2* cells grew slightly better than when the cells were maintained in CL (**Fig. 8A**), indicating that the loss of CreTPT3 might be partially compensated for by starch turnover during the night (or allow for some repair of cellular damage that might accumulate during the day), but to a lesser extent than in plants. We observed night-time starch degradation in *tpt3* cells, although immediately following the night period a higher level (∼5-fold) of undegraded starch remained relative to that of WT cultures (**Fig. 8C**). These data suggest that starch mobilization may partially compensate for a CreTPT3 deficiency in Chlamydomonas.

**Fig. 8:**
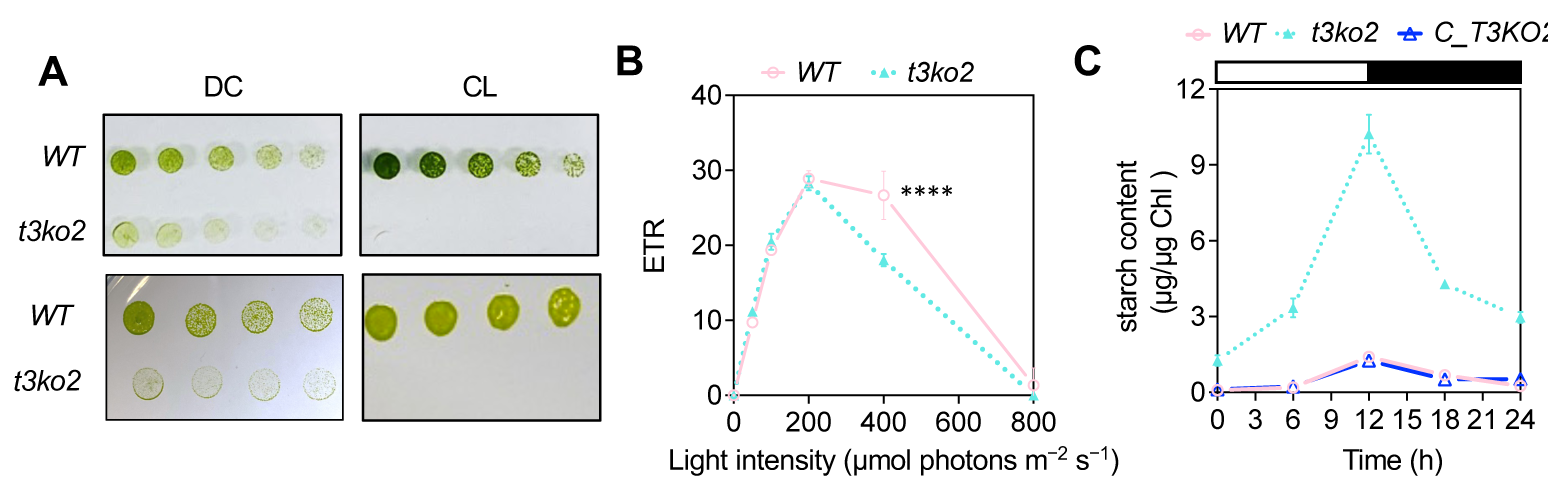
Effects of deletion of CreTPT3 over diurnal cycle. (**A**) Growth of WT, and *t3ko2* strains on TP agar plates for 4 days either under diurnal cycle (DC) (left panel; light : dark/ 12h : 12h) or continuous light (CL) (right panel) at an intensity of 60 μmol photon m^-2^s^-1^ (upper panel) or HL (lower panel: 450 μmol photon m^-2^s^-1^). The dilution series are 3, 1.5, 0.75, 0.375 μg/mL chlorophyll. (**B**) Photosynthetic electron transport rate (ETR) of cells grown under a diurnal rythym with the light period at an intensity of 60 μmol photon m^-2^s^-1^. (**C**) Accumulation of starch during growth under a diurnal rythym at a light intensity of 60 μmol photon m^-2^s^-1^ .

The mechanism by which starch breakdown products are exported from the chloroplast during the night in Chlamydomonas remains largely unknown, although it could involve various transporters including MEX1 (like in plants). Chlamydomonas MEX1 can transport starch break down products in the form of glucose and/or glucose phosphate, but there is no evidence suggesting that it can export maltose since a *mex1* mutant did not accumulate maltose or exhibit growth impairment (Findinier et al., 2017). We speculate that the loss of TPT3 might be partially compensated for by starch turnover in the dark, with degradation products exported via MEX1 or other pPTs that are highly expressed in the dark and downregulated in the light, such as CreTPT10 and CGL51 (**Supplementary Fig. 1D**). However, even if hexose-P is exported from the chloroplast, it may not be readily converted to triose-P (see below), which would fuel the TCA cycle and respiration and serve to support anabolic processes.

### *t3ko2* experiences oxidative stress

In plants, the export of sugars and other molecules (e.g. redox equivalents, ROS) can serve as signals that coordinate chloroplast and nuclear gene expression during acclimation to HL (Häusler et al., 2014; Zirngibl et al., 2022). We probed the impact of impaired triose-P export in *t3ko2* on ROS production and accumulation in various subcellular compartments following exposure of the cells to ML or HL. In plant cells, H_2_O_2_ is produced as a side-product of cellular processes including PET, mitochondrial respiration, and substrate level oxidation (Smirnoff and Arnaud, 2019; Foyer and Noctor, 2016). While ROS stability is generally low, it can accumulate in cells experiencing oxidative stress, with H_2_O_2_ being the most prevalent species that can function as a redox messenger (Li and Kim, 2022). Moreover, the trafficking of H_2_O_2_ into or out of different cellular compartments can trigger activation of other retrograde and anterograde signals that may coordinate activities among the compartments, including the nucleus (Exposito-Rodriguez et al., 2017; Mittler et al., 2022; Shapiguzov et al., 2012). Upon exposure to HL, the *t3ko2* mutant accumulated more stromal H_2_O_2_ than WT cells. The cytosolic H_2_O_2_ levels in the mutant exhibited a similar increase, which may reflect the ability of this metabolite to rapidly diffuse from the chloroplast and into the cytoplasm (**Fig. 6B, C**). Furthermore, it is unlikely that the mitochondrion contributes to an increase in H_2_O_2_ in *t3ko2* since no (or little) increase in accumulation of intramitochondrial H_2_O_2_ was observed in the mutant in either LL or ML (**Fig. 6D**). An increase in H_2_O_2_ accumulation in the nucleus of *t3ko2* relative to WT cells was also observed, although the light intensity (HL or LL) did not alter these levels in either WT or the mutant (**Fig. 6E**); a previous report showed that the nuclear H_2_O_2_ level was not significantly affected in WT Chlamydomonas cells following a HL exposure (Niemeyer et al., 2021), which may reflect both the level of H_2_O_2_ accumulation and barriers that limit its diffusion. The higher levels of H_2_O_2_ in the nucleus of *t3ko2* cells may trigger retrograde signals that modulate nuclear gene expressions, which in turn could ameliorate some of the negative effects of ROS and elicit repair of any damage experienced by the photosynthetic apparatus. A similar response may be elicited in WT cells at higher intensity actinic light.

Superoxide is predominantly produced by PSI, which can enzymatically be converted into H_2_O_2_ (Foyer, 2018). Highly reduced PSI in the *tpt3* mutant can lead to an increase in the production of H_2_O_2_. It was also shown that H_2_O_2_ can be synthesized in thylakoid membranes as a consequence of the oxidation of plastoquinol (PQH_2_), suggesting a positive correlation between the redox state of the PQ pool and the generation of H_2_O_2_ (Khorobrykh et al., 2015). A similar finding was noted for both *Nicotiana benthamiana* and Chlamydomonas based on the use of hypersensitive H_2_O_2_ sensors (Exposito-Rodriguez et al., 2017; Niemeyer et al., 2021). These two organisms were shown to accumulate more stromal H_2_O_2_ in HL, which was dependent on photosynthesis. We observed this positive correlation between PQ pool reduction and the accumulation of H_2_O_2_ in the *t3ko2* mutant; the PQ pool was more reduced in *t3ko2* relative to the WT cells after exposing the cells to 105 s of HL (**Fig. 4D)**. In parallel, there was a marked increase in stromal H_2_O_2_ following 2.5 min of HL (**Fig. 6B**). Therefore, hyper-reduction of the PQ pool in the mutant likely results in elevated stromal H_2_O_2_ accumulation, suggesting that CreTPT3 activity and the export of triose-P from the chloroplast is critical for maintaining low level synthesis/accumulation H_2_O_2_ and sustaining a high rate of PET in HL; the export of fixed carbon relieves the redox pressure and lessens ROS formation. Additionally, Cre*TPT3* is the most nutrient-deprivation responsive/upregulated of the *pPT* family genes; it responds strongly to nitrogen, sulfur, and iron deprivation (**Supplementary Fig. 9 B-D**). These findings are in accord with the hypothesis that the ability to traffic fixed carbon from the chloroplast is important for both the distributing carbon to other cellular compartments and relieving oxidative stress in the organelle.

### CreTPT3 is critical for maintaining intracellular partitioning of fixed carbon

Why is the phenotype of the Chlamydomonas *tpt3* mutant so severe? Land plants contain the entire glycolytic pathway in both the chloroplast and cytosol while the pathway is partitioned between two compartments in Chlamydomonas; 90% of the upper activities of the pathways (from F6Pà 3PG) was associated with the plastid while over 95% of the activities of the lower part of the pathway (3PGàPyruvate) occurred in the cytosol (Klein, 1986; Rochaix et al., 1998). The oxidative pentose phosphate pathway also appears to be in the chloroplast (Klein, 1986). The partitioning of glycolysis between the chloroplast and cytosol is supported by the comparative quantification of the metabolites, with glucose-1-P, fructose-6-P, and fructose-1,6-P2 being exclusively in the chloroplast and 2-phosphoglycerate only in the cytosol (Klöck and Kreuzberg, 1991). Recently, it was suggested that the flux of metabolites through hexose-P is negligible in the Chlamydomonas cytosol (Treves et al., 2022), possibly because of the absence of glycolytic reactions that would facilitate its metabolism. Therefore, even if hexose-P is exported from the chloroplast, it would likely not be rapidly metabolized or maintain rapid cell growth. Overall, the results strongly suggest that triose-P exported from the Chlamydomonas chloroplast is likely the major source of fixed carbon transported into the cytoplasm of the cell, facilitating algal growth in the light.

Thus, we hypothesize that the export of triose-P would drive the cytosolic segment of glycolysis and downstream metabolic pathways. This hypothesis is supported by the metabolite analysis; specifically, a marked increase of most metabolites associated with the upper-glycolytic/gluconeogenic pathways, and a significant decrease of some metabolites of the TCA cycle upon exposure of the mutant cells to either LL or ML (**Fig. 3E-G, Supplementary Fig. 8B, C**). In contrast, elimination of the chloroplast-targeted TPT1 protein of Arabidopsis showed no significant phenotype, although growth was retarded in the TPT/XPT double mutants (Hilgers et al., 2018a). Furthermore, based on Pearson correlation analyses presented in **Supplementary Fig. 11** and **Table. 4**, Cre*TPT3* is co-expressed with many genes involved in respiratory electron transport and the major ATP transporters located on the mitochondria and chloroplast envelope membranes. In addition, transcript levels of some genes involved in starch degradation, glycolysis, the TCA cycle, and malate/OAA shuttle shared a high correlation coefficient with Cre*TPT3*. Together, these data indicate that the export of triose-P from the chloroplast is closely linked to central energy metabolism in Chlamydomonas, starting with the production of triose-P in the chloroplast by the CBBC or starch degradation (chloroplast localized reactions of glycolysis), followed by transport to the cytosol which houses the remaining reactions of glycolysis. The products of glycolysis can be trafficked to the mitochondria where they can be used to drive the TCA cycle, respiratory metabolism, and the generation of ATPs (**Supplementary Fig. 11**). Hence, triose-P is the major photoassimilate routed from the chloroplast, supplying substrates for downstream metabolic processes.

## Summary

As depicted in **Fig. 7**, we propose that various tiers of regulation are responsible for the physiological responses of the Chlamydomonas *t3ko2* mutant. When *t3ko2* cells are transferred from LL to ML, the triose-P pool and metabolites derived from that pool accumulate because of the reduced capacity of the strain to move triose-P out of the chloroplast where it could be further metabolized. Some compensation may occur through the activity of other transporters, although expression of Cre*TPT2* is especially low during the day (in the light) and the transport of hexose phosphate may not compensate for the loss of CreTPT3 because the cytoplasm does not have (or has little of) the activities of glycolysis that would convert hexose-P to DHAP. The compromised ability to export fixed carbon from the chloroplast also suppresses CBBC activity, causes hyper-reduction of PET and the accumulation of ROS (which would inhibit photosynthetic activity). The highly diminished export of triose-P to the cytoplasm would compromise respiration and downstream biosynthetic processes. Furthermore, hyper-reduction of PET and accumulation of carbon metabolites in the stroma would activate AGPase through allosteric regulation and by the FDX/TRX (ferredoxin/thioredoxin) and NTRC (NADPH-dependent thioredoxin reductase C) redox systems, which would result in starch hyper-accumulation (Ballicora et al., 2000; Lepistö et al., 2013) (**Fig. 7**). Increased ROS accumulation in the mutant chloroplasts and an elevated PET redox state would also elicit the generation of retrograde signals that mediate changes in nuclear gene expression (Wakao and Niyogi, 2021; Suzuki et al., 2012; Shapiguzov et al., 2012), stimulating the synthesis of specific activities that may function to ameliorate the impact of the hyper-reduced state attained in the chloroplast.

## Material and methods

### Strains and culture conditions

WT Chlamydomonas M10 (CC-4403, isogenic line derived from CC-124) was used as the parental strain for the generation of knockout mutants. Cultures were routinely cultivated in growth chambers (LED-41L2, Percival Scientific, Inc.) at 25°C with continuous shaking on an orbital shaker (VWR OS-500 Shaker) at 120 rpm, in Tris-Acetate-Phosphate (TAP) medium (Harris, 2009). Cultures were illuminated with continuous cool white LEDs (LED-41L2, Percival Scientific, Inc.) at low light (LL, 30 μmol photons m^-^_2_ s^-1^). Experiments were mostly performed with cells grown in the photoautotrophic TP medium (TAP medium without acetate) and in some cases, in TAP medium, at 25°C, and sparged with air while being shaken at 120 rpm in a growth chamber (LED-41L2, Percival Scientific, Inc.). For growth assays, cultures were inoculated to a density of 0.02 at OD750 nm (∼1×10^5^ cells/mL) in TP sparged with air under either LL or moderate light (ML, 250-300 μmol photons m^-^_2_ s^-1^) intensities. For spectrophotometric and chlorophyll fluorescence analyses, the experimental design is described in **Supplementary Fig. 5**. Growth assays on solid medium were performed with cultures spotted onto the medium at different dilutions (indicated in the text) and exposed to different light intensities; spot tests for photoautotrophic growth were on TP agar plates and for mixotrophic growth on TAP agar plates. Agar plates were incubated for 7 d under either LL or ML (cool white LED) at 25°C.

### Reconstitution into liposomes and transport assays

The procedures for the construction and expression of Cre*TPT2* and Cre*TPT3* in yeast are described in supplemental information (SI). For uptake studies, yeast membranes from cells with and without recombinant His-tagged CreTPT2 or CreTPT3 were enriched and reconstituted into 3% (w/v) L-alpha-phosphatidylcholine using a freeze-thaw-sonication procedure (Linka et al., 2008). The reconstituted liposomes were preloaded with 30 mM Pi or phosphorylated metabolites to be tested as potential transport substrates. As a negative control for antiport activity, liposomes were also generated without metabolite preloading. The external counter-exchange substrate was removed via gel filtration on Sephadex G-25M columns (GE Healthcare). Transport assays were initiated by adding 0.25 mM [alpha-^3^_2_P]-phosphoric acid (6,000 Ci/mmol) to the medium bathing the liposomes and performed as previously described by Linka et al. (2008). Measurements of Km for Pi and competitive inhibition constants (Ki) are described in supplemental information (SI).

### Vector construction, transformation, and subcellular localization

The pRam118_VENUS plasmid, which harbors the VENUS gene and the *AphVII* cassette (confers resistance to hygromycin) (Cre01.g045550)(Kaye et al., 2019), was used to express Cre*TPT2* (Cre06.g263850_4532) and Cre*TPT3* (Cre01.g045550_4532). The step-by-step description is in SI. 2-4 μg of the engineered plasmids was linearized and added to 250 µL of a cell suspension of ∼3×10^8^ cells/mL per reaction. The GeneArt MAX Efficiency Transformation Reagent for algae (Invitrogen) was used for introducing the plasmid into the algal cells by electroporation according to the instructions provided by the manufacturer. Transformants were selected on solid TAP medium containing hygromycin (10 μg/mL; Enzo Life).

Drug resistant transformants were visualized for VENUS fluorescence as previously described (Kaye et al., 2019). In brief, transgenic cell lines resistant to hygromycin were screened for VENUS fluorescence using a microplate reader (Infinite M1000; TECAN). Excitation and emission settings were: VENUS, excitation at 515 nm, bandwidth 12 nm and emission at 550 nm, bandwidth 12 nm; chlorophyll excitation was at 440 nm, bandwidth 9 nm and emission was at 680 nm, bandwidth 20 nm. The TCS SP8 confocal laser-scanning microscope (Leica) was used to visualize the VENUS fluorescence signal (Kaye et al., 2019).

### CRISPR-CAS9 mediated mutagenesis

The Chlamydomonas WT strain CC-124 was used for mutant generation. WT cells were cultured under CL (50 μmol photons m^-^_2_ s^-1^) for 2 d to a density of 3-5×10^6^ cells/mL. The cells were then concentrated to 2×10^8^ cells/mL in 0.5×TAP medium supplemented with 80 mM sucrose. Two single guide RNAs (sgRNA) were designed by CHOPCHOP and synthesized by Integrated DNA Technologies (IDT). The sequences of the generated sgRNAs are: *TPT2*-sg (5′-AUAAGGGCAAGGACAUGUCAGGG -3) for editing exon 8, *TPT3*-sg1 (5′-CGCUGGGCGTCACUUCCCGGCGG -3′) for editing exon 1, and *TPT3*-sg2 (5′-AAGGCCGCUAUCGCCAACGUGGG -3′) for editing of exon 7. The protocol for disruption of Cre*TPT2* and Cre*TPT3* was adapted from (Findinier et al., 2019) and is described in SI.

### Complementation of mutants *tpt3* mutants

Mutant strains were transformed with the linearized pRam118_CreTPT3 plasmid. Transformed cells were selected in ML and screened for VENUS fluorescence. Colonies exhibiting VENUS fluorescence and an *AphVII* cassette knock-in at the CAS9 target site were examined for accumulation of the CreTPT3 protein by immunodetection using CreTPT3 antibodies generated by GenScript USA Inc (Piscataway, USA). Immuno-positive colonies were subjected to growth assays using spot tests under ML on either solid TAP or TP medium.

### P700 activity measurements

Absorbance spectroscopy [JTS-100 spectrophotometer (SpectroLogiX, TN)] to measure P700 activity was performed with dark-adapted liquid cultures (15 μg/mL chlorophyll, in 20 mM HEPES-KOH, pH 7.2, and 10% ficoll) as previously described (Clowez et al., 2021). Actinic light was provided by an orange LED (165 μmol photons m^-^_2_ s^-1^) for PSI oxidation, followed by a saturating pulse and dark incubation. 20 μM DCMU and 1 mM hydroxylamine were added to the cell suspension to inhibit linear electron flow (LEF) prior to the measurement. P700 activity was measured by monitoring the absorbance at 705 nm (interference filter 6 nm FWHM was used to create a narrow excitation beam); the absorbance at 740 nm was used to correct for an unspecific contribution to the 705 nm signal.

### Chlorophyll fluorescence analysis

Chlorophyll fluorescence used to evaluate photosynthetic electron transport was monitored with a DUAL PAM-100 fluorometer. Cells were acclimated in the dark for 20 min prior to illumination at increasing light intensities (0, 10, 50, 100, 200, 400, 800, 1000, 1200 μmol photons m^-^_2_ s^-1^) for 2 min at each intensity, or at a constant intensity of 450 μmol photons m^-^_2_ s^-1^ for 10 min to evaluate 1-qL. 1 mM CO_2_ (NaHCO3) was added to the reaction mix as an electron acceptor for the CBBC.

### ROS measurements and roGFP2 imaging analysis

All strains were grown photoautotrophically in LL for 16-24 h. After dilution with fresh medium, the cultures were transferred to ML and stained with CM-H2DCFDA (Thermo Fisher Scientific) as described in (Kong et al., 2018) for detecting ROS. A step-by-step protocol is given in SI.

Constructs containing chloroplast or mitochondria targeting sequences (Crozet et al., 2018) fused to codon-optimized roGFP2 (Vevea et al., 2013) were transformed into WT and mutant strains. Transgenic cell lines were screened for green roGFP2 fluorescence using a microplate reader (Infinite M1000; TECAN); excitation was at 488 nm, bandwidth 9 nm and emission at 525 nm, bandwidth 10 nm. Cells with strong green fluorescence were cultured as depicted in **Supplementary Fig. 5**. Signals from the transformed lines were visualized using a TCS SP8 confocal laser-scanning microscope (Leica). roGFP2 signals were collected and analyzed as previously described (Vevea et al., 2013). Using the sequential setup of the SP8, roGFP2 signals were collected at the emission wavelength of 510-550 nm immediately following excitation at 405 nm and 488 nm (Vevea et al., 2013). The degree of roGFP2 oxidation was analyzed as the ratio of the emission signals after excitation at 405 nm and 488 nm (Vevea et al., 2013).

Plasmids harboring H_2_O_2_ roGFP2-Tsa2ΔCR sensors (Niemeyer et al., 2021) were obtained from the Chlamydomonas Resource Center (https://www.chlamycollection.org/) (**Supplementary Table. 3)**. Cells in which roGFP2-Tsa2ΔCR was targeted to the stroma, cytosol, mitochondrial matrix, or nucleus were initially grown in LL in TAP to exponential growth phase. Cells were then grown in TP in LL for 2 h and shifted either to HL or very low light (10 µmol photons m^−^_2_ s^−1^). Prior to collecting the algal samples, N-Ethylmaleimide (NEM) (in 100-mM MES–Tris buffer, pH 7.0) was added to a centrifuge tube to trap the oxidation state of the sensor. The final concentration of NEM is 10 mM after adding the algae culture. After centrifugation (1459 x g, 3 min), cells were resuspended in 100-mM MES–Tris buffer (pH 7.0) to reach the chlorophyll concentration of 30 µg/mL; and roGFP2 fluorescence was measured in a plate reader (Infinite M1000; TECAN). For calibration and data calculation, fully oxidized sensors in the control samples were prepared by adding H_2_O_2_ to a final concentration of 5 mM, and fully reduced sensors were prepared by adding DTT to a final concentration of 100 mM. Signals were detected using the excitation wavelengths of 410 and 488 nm and the emission wavelength of 514 nm. The degree of sensor oxidation (OxD) was calculated using the previous equation as described in (Niemeyer et al., 2021).

### Photosynthesis-Irradiance curve

Photosynthesis-Irradiance curves were measured using a custom Pt-Ag/AgCl polarographic electrode system (ALGI) with a water jacketed (for temperature-control), 1 mL glass reaction chamber. A step-by-step protocol is provided in SI.

### Starch and TAG quantification

Cells were grown under LL to mid-exponential phase, diluted to 0.5 μg/mL chl with fresh medium and transferred to and then grown in ML for 24 h or more, as indicated. Starch was measured according to (Klein and Betz, 1978), with slight modification (see SI). Triacylglycerol (TAG) levels were estimated by a fluorometric assay using the dye Nile Red (Thermo Fisher)(Yu et al., 2009). A Nile Red solution (500 μg/mL in acetone) was added to 1 mL of the cell suspensions to a final concentration of 0.5 μg/mL. Samples were then incubated at room temperature for 30 min, and the Nile Red fluorescence emission quantified at 575 nm following excitation at 530 nm using a microplate reader (Infinite M1000; TECAN).

### Metabolic analysis

Cells grown in LL were shifted to ML for 45 min or 6 h, as indicated. 45 mL of culture was rapidly quenched in cold saline solution (-2 to -3°C), extracted using cold methanol and then analyzed by LC-MS/MS. Quenching and analysis of metabolites were performed, with modifications, according to (Sake et al., 2020). A step-by-step protocol is described in SI. Metabolite extracts were analyzed using LC-MS/MS, as adapted from (Young et al., 2011). A Phenomenex 150 mm x 2 mm Synergi Hydro-RP column was used on an Agilent 1200 Series HPLC system coupled to an AB Sciex 5500 QTRAP system. LC was performed with a liquid injection volume of 20 μL and a gradient elution with 10 mM tributylamine and 15 mM acetic acid (aqueous phase) in acetonitrile (organic phase) (reagent B) at a constant flow rate of 0.3 mL/min, and a constant temperature of 40°C. The gradient profile of the organic phase is as follows: 0% B (0 min), 8% B (10 min), 16% B (15 min), 30% B (16.5 min), 30% B (19 min), 90% B (21.5 min), 90% B (28 min), 0% B (28.5 min), and 0% B (35 min). MS analysis was performed in negative mode using a multiple reaction monitoring (MRM) acquisition method. Data acquisition was performed on the ABSciex Analyst 1.7 software. Absolute quantification of intracellular metabolites was performed using the quantitation mode on the Analyst software. All chemicals used for metabolite extraction and LC-MS/MS analysis were Optima grade reagents.

## Acknowledgments

WH thanks Dr. Masayuki Onishi for advice on the generation of CRISPR mutants and Adrien Burlacot for help with PQ pool measurements.

## Author contributions

WH, AK, ARG and MP conceptualized the study. WH generated the CRISPR mutants as well as the complemented strain, localized the TPT2 and 3 proteins, analyzed the photosynthetic performances and starch changes over diel cycle. AK performed the experiments of photosynthetic O_2_ evolution, extraction of samples for LC-MS/MS, data analysis of LC-MS/MS. AP and NL performed the reconstitution into liposomes and transport activity assays. MM and NB performed LC-MS/MS and analyses of metabolite data. WH and YW analyzed the redox status in the cell. BR performed data mining for transcriptome and the Nile red staining. JF and NF analyzed starch changes upon transition from LL to ML. WH, PR and ARG analyzed PSI and PSII activities. JF and BR constructed roGFP2 associated vectors. WH and ARG wrote the manuscript. ARG, MP, AK, NL, NB, AP, MM and all the other authors helped writing and revising the manuscript.

## Distribution of materials

All manuscripts must include the following statement as an unnumbered footnote: "The author(s) responsible for distribution of materials integral to the findings presented in this article in accordance with the policy described in the Instructions for Authors (https://academic.oup.com/plcell/pages/General-Instructions) are: Weichao Huang and Arthur Grossman.

## Supplementary Materials

### Materials and methods

#### Expression of CreTPT2 and CreTPT3 in yeast

The DNA sequence encoding the mature CreTPT2 or CreTPT33 protein was codon-optimized for expression in *Saccharomyces cerevisiae* (GeneART, ThermoFisher Scientific). This coding sequence was inserted in frame with an N-terminal His tag into the yeast vector pYES-NTa (ThermoFisher Scientific) using Gibson cloning (NEB). Briefly, pYES-NTa was linearized with BamHI and the Cre*TPT2* and *TPT3* cDNA amplified with the primer pairs: fwd:5’-gacgataaggtacctagcGCTGCTGCTGTTCCAGCTG-3’ and rev: 5’-agaattccaccacactgTTAAGCAGCTTCTGGCTTAACATC-3’ (or Cre*TPT2*); fwd: 5’ - gacgataaggtacctagcGCTTCTGCTGCTGATGCTC-3’ and rev: 5’ - agaattccaccacactgTTAAGCGGCAGCTGGAGAAG-3’ (for Cre*TPT3*). This cDNA was ligated into the linearized pYES-NTa vector, which was then transformed into the yeast strain INVSc1 (MATa, his3D1, leu2, trp1-289, ura3-52 / MATa, his3D1, leu2, trp1-289, and ura3-52, Thermo Fisher Scientific) using the lithium-acetate/PEG method (Gietz and Schiestl, 2007). Transformed yeast cells were selected on synthetic complete medium containing 2% (w/v) glucose with the uracil auxotrophic marker. Galactose-inducible expression of His-CreTPT2 or CreTPT3 in yeast was performed as described in (Linka et al., 2008). The presence of the His-tagged fusion protein was verified by standard SDS-PAGE and immunoblot analysis using an anti-His-antibody conjugated with horseradish peroxidase (Miltenyi Biotech).

#### Transport activity assays

The Km for Pi was determined by measuring the initial velocity at each of six external Pi concentrations between 0.05 and 5 mM. To obtain competitive inhibition constants (Ki), the uptake of 0.25 mM Pi into liposomes containing 30 mM Pi was measured over a 4 min period in the presence of increasing external competitor–concentrations (0.05 - 5 mM). Three biological replicates were performed for all described experiments. GraphPad Prism software version 9.3.0 was used for non-linear regression analyses of the kinetic data.

#### Vector construction

The plasmid of pRam118_VENUS was linearized with HpaI (NEB). Primers: g*TPT2*_pRam118_f, g*TPT2*_pRam118_r and g*TPT3*_pRam118_f, g*TPT3*_pRam118_r (**Supplementary Table 1**) were used to amplify Cre*TPT2* and Cre*TPT3* genomic DNAs containing an overlap with the linearized pRam118_VENUS vector. For generating the plasmids pRam118-CreTPT2&VENUS and pRam118-CreTPT3&VENUS, genomic DNA of Cre*TPT2* or Cre*TPT3* was assembled with pRam118_VENUS plasmid using Gibson assembly (Gibson et al., 2009). The plasmid of pRam118-CreTPT2&VENUS (2-4 μg) linearized with AseI (NEB), was transferred into the Chlamydomonas M10 strain by electroporation.

A 1,000 bp sequence upstream of Cre*TPT3* and containing the promoter region of the gene was amplified using the primers TPT3pro1000_f and TPT3pro1000_r (**Supplementary Table 2**). The 3’ UTR of Cre*TPT3* was amplified using TPT_3UTR_f and TPT3_3UTR_r (**Supplementary Table 2**). The *PSAD* promoter and the *RBCS2* 3’UTR of pRam118-CreTPT3&VENUS were replaced by the amplified fragments of the 1,000 bp upstream region and the 3’ UTR of Cre*TPT3*, respectively. This final vector, designated pRam118_CreTPT3, contains the original Cre*TPT3* promoter (driving expression of Cre*TPT3*), the genomic DNA sequence of Cre*TPT3*, VENUS and the Cre*TPT3* 3’ UTR plus the *AphVII* cassette. To locate CreTPT3 and complement the *tpt3* mutant, the mutant was transformed by electroporation with a total amount of 2-4 μg (∼500 ng/µL) of pRam118_CreTPT3 that was linearized with AseI.

#### CRISPR-Cas9 mediated mutagenesis

Prior to electroporation, Cas9 (IDT) and sgRNAs were incubated together at 37 °C for 30 min. Approximately 500 ng PCR product of the *AphVII* cassette, which confers resistance to hygromycin, was added to the RNP (ribonucleoprotein) mixture. 250 μl aliquots were electroporated using Super Electroporator NEPA21 type II (NEPA GENE). After 16 h of recovery in very low light (10-15 μmol photons m^-^_2_ s^-1^), cells were plated onto solid TAP medium containing 10 μg/mL hygromycin. Sense or antisense-oriented knock-ins of *AphVII* were determined by amplification using primer pairs with one primer annealing to the genomic sequence and the other to the inserted sequence (**Supplementary Table 2**). The amplified fragments were sequenced to verify the insertion sites (ELIM BIOPHARM, Hayward, USA).

#### ROS measurements and roGFP2 imaging analysis

Briefly, after various treatment (e.g. 48 h in ML), 10 million Chlamydomonas cells were pelleted and washed once with 1×PBS. The cells were then resuspended in 1×PBS containing 8 μM CM-H2DCFDA and incubated at room temperature in the dark for 30 min. Following this incubation, the cells were washed three times with 1×PBS buffer, the fluorescent signals were either visualized using TCS SP8 confocal laser-scanning microscope (Leica) or quantified with a microplate reader (Infinite M1000; TECAN). Excitation and emission settings for the microscope were: 488 nm/510-530 nm HyD SMD hybrid detector for reactive oxygen species (ROS), and 488 nm/650-700 nm-HyD SMD hybrid detector for chlorophyll autofluorescence. Excitation and emission settings for the plate reader were as follows: ROS excitation 488/5 nm and emission 530/12 nm; chlorophyll excitation 514/5 nm and emission 690/5 nm.

#### Starch analysis

In brief, cells were collected by centrifugation and pigments extracted with methanol. Dried pellets were resuspended in water and heated at 100°C to break the cells and release the starch. After cooling, amyloglucosidase and α-amylase (2.25 U/mL) were used to hydrolyze the starch and the products of hydrolysis were quantified using a Glucose Colorimetric Detection Kit (Thermo Fisher).

#### Photosynthesis-Irradiance curve

The YSI 5331A electrodes (Yellow Springs Instruments) were polarized at −0.8 V. The cultures used for the assays were concentrated to 2.5-5 µg mL^-1^ chlorophyll, supplemented with 15 µL of 0.5 M sodium bicarbonate in water and then purged with 1% CO_2_/99% He. Using a gas tight syringe, the sample was transferred into the reaction chamber that was also purged with 1% CO_2_/99% He. The rate of change in O_2_ levels was measured sequentially at the light intensities 100, 200, 600, 1000 µmol photons m^-^_2_ s^-1^ (photosynthetic active radiation, PAR); each intensity was maintained for 3-5 min followed by a 3-min intervening dark period, and then the light level was raised to the next higher intensity (stepped change) until the full range of intensities was tested. At 1,000 µmol photons m^-^_2_ s^-1^, the light was held for 10 min followed by an 8-min dark period. Before the measurement of each experimental series, the electrodes were calibrated with air (∼21% O_2_) and 1% CO_2_/99% He mixture (0% O_2_). The initial slope of the response was used to determine the O_2_ evolution rate.

#### Extraction of samples for LC-MS/MS

The quenching solution, filtered saline (9 g/L NaCl), was prechilled to 4°C in a refrigerator. 30 mL of the quenching solution was transferred to a 50 mL conical tube kept in an ice bath mixed with salt to depress the temperature to between −3 °C and −1 °C. 15 mL of the culture was rapidly plunged into the 30 mL of the quenching solution and the samples were centrifuged at 4,000 rpm for 10 min at 4 °C. Cell pellets were washed with fresh prechilled saline solution and centrifuged again at 4,000 rpm for 10 min in a 2 mL centrifuge tube. For each replicate, 45 mL of the culture was sampled and the cell pellets were pooled. Washed pellets were frozen at −80 °C until extraction. Metabolite extraction and further analysis were modified from (Young et al., 2011). To each cell pellet, 500 μL methanol was added along with the internal standards ribitol and PIPES to a final concentration of 2 µM each. Samples were then vortexed for 30 sec, frozen in liquid nitrogen as described in (Winder et al., 2008), and allowed to thaw at 0 °C. This vortex-freeze-thaw cycle was repeated twice more before the samples were centrifuged at 10,000 x g at 1 °C for 5 min. The supernatant was collected, and the remaining pellets were extracted twice more with 500 μL of a 50:50 mixture of methanol and water, with 3 vortex-freeze-thaw cycles done for each extraction. Supernatants from each extraction procedure were pooled and dried on a Thermo Fisher SpeedVac Concentrator. Dried extracts were then resuspended in 500 μL water and cleaned to remove any residual large cell debris by filtration, first through a 0.22 μm pore size Spin-X centrifugal tube filter, followed by 10 K molecular mass cutoff filters, followed finally by 3 K molecular mass cutoff filters. After each filtration step, filters were rinsed with 50 μL water, which was added to the total sample volume for subsequent steps. Filtered extracts were dried again and resuspended in 200 μL water for LC-MS/MS analysis.

#### Protein extraction and immunoblot analysis

Affinity-purified polyclonal antibodies to CreTPT3 were custom-made by GenScript. The antigen sequence used for antibody generation was KSWSFGRPVTKQEF. Chlamydomonas cells were grown in liquid cultures to 2–5 x10^6^ cells ml^-1^ and collected by centrifugation (1459 x g, 5 min). Cells were resuspended in resuspension buffer [(5 mM HEPES, pH 7.5, KOH), 10 mM EDTA (pH 7.5, NaOH), 1x protease inhibitor “complete EDTA-free” (Roche)]. For the disruption of cells, bead-beating was performed using a mini beadbeater (Biospec) in two cycles of 30 sec min^-1^, with a 1 min period of cooling on ice between cycles. Disrupted cells were centrifuged at 4 °C, 30 min, 14000 x g in a microfuge. The supernatant was removed, the pellet was resuspended in sample buffer (resuspension buffer supplemented with 100 mM Na_2_CO3, 100 mM DTT, 2% WT/vol SDS, and 12% WT /vol sucrose).

SDS-PAGE was performed using a 12% polyacrylamide gel (Bio-Rad), with the electrophoresis for 90 min at 120 V. The proteins within the gel were transferred onto a PVDF membrane (Bio-Rad) using Trans-Blot Turbo (Bio-Rad). The detection was performed using Clarity Max Western ECL Substrate (Bio-Rad).

#### RT-qPCR

RT-qPCR, was performed as described previously (Kaye et al., 2019). Briefly, total RNA was isolated using the RNeasy Plant Mini Kit (Qiagen) and treated with DNase I (Qiagen). First-strand cDNA was generated by reverse transcription of 0.5 µg total RNA using the iScript cDNA Synthesis Kit (Bio-Rad). Real-time PCR using the Roche Light Cycler 480 was performed with the SensiMix NO-Rox SYBR Green I Kit as described by the manufacturer (Bioline). Oligos used in this research for analyzing the expression levels of Cre*TPT10/TPT2/TPT3/CGL51* are listed in **Supplemental Table 1**. *mRNA levels were calculated using DeltaC (T) method, CBLP gene was used as an internal control*.

#### Analysis of transcript levels in response to different conditions

In order to analyze the transcript level of Cre*TPT* genes under different conditions, we obtained RNAseq data published previously in which cells experienced a diurnal cycle, nitrogen starvation, sulfur starvation and Fe_2_^+^ starvation (Strenkert et al., 2019; Zones et al., 2015; Ngan et al., 2015; González-Ballester et al., 2010; Urzica et al., 2013). The log2 fold change of each gene was calculated based on the ratio of transcript abundance at different time point over transcript level at time point 0 h. The fold change was visualized by a heatmap and the color bar is the scale of log2 (fold-change).

#### Correlation analysis based on Cre*TPT3* transcription abundance under different conditions

The Z score of each gene was calculated as follows: the mean transcript level of each gene from all of the conditions was subtracted from each gene’s transcript level. This difference was divided by the standard deviation of each gene’s transcript level under all conditions to get Z score numeric values. The whole Z score matrix for every gene under all conditions was used to calculate the Pearson correlation matrix by using python. Dataframe.corr() function. The genes highly correlated with TPT3 were selected if their correlation value is higher than 0.75. The Pearson correlation value of TPT3-highly-correlated genes were plotted based on their KEGG pathway.

#### In vivo polymeric carbohydrate staining

Lugol’s iodine (aqueous solution of 1.8% iodine and 3.0% potassium iodide) was used to stain in vivo polymeric carbohydrate. Briefly, 0.5 mL of the cell culture was pelleted, resuspended in 10 µL of Lugol’s iodine and visualized under a bright-field microscope. Lugol’s iodine selectively binds to alpha-1,4 glucans and stains it blue-black.

## Supplementary Figures

**Supplementary Fig. 1:**
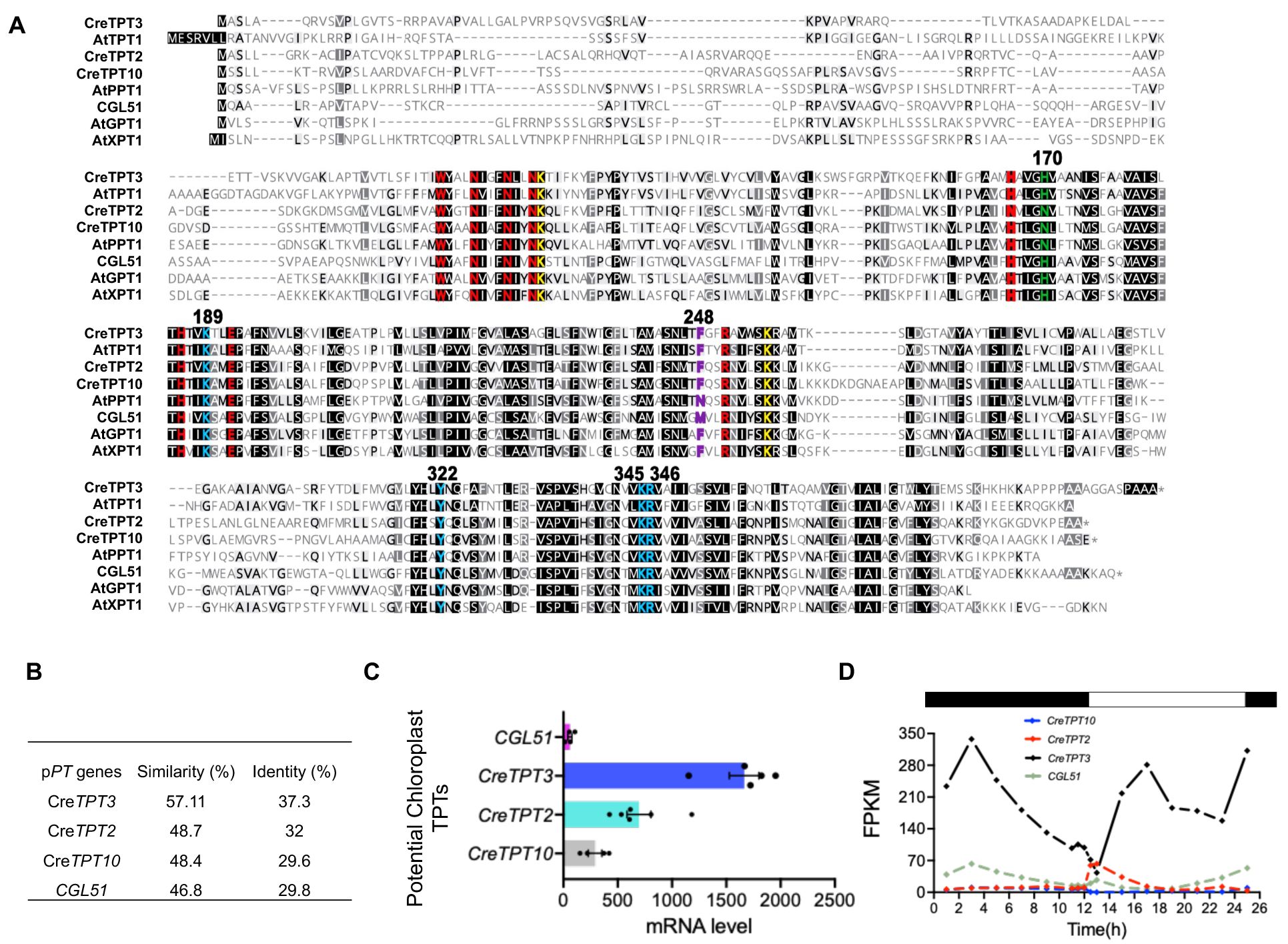
Features of putative plastid phosphate transporter (pPT) in *Arabidopsis thaliana* and *Chlamydomonas reinhardtii.* **(A)** Alignment of sequences of pPTs in Arabidopsis and Chlamydomonas. Plastid putative sugar phosphate transporters from Chlamydomonas are CreTPT10 (Cre08.g379350_4532), CreTPT2 (Cre06.g263850_4532), CreTPT3 (Cre01.g045550_4532), and CGL51 (Cre16.g663800_4532). Sequences from *Arabidopsis thaliana* are AtTPT1 (AT5G46110.1), AtPPT1 (AT5G33320.1), AtGPT1 (AT5G54800.1) and AtXPT1 (AT5G17630.1). Conserved substrate binding pockets are colored red. Phosphate-binding residues are colored blue. The gate capping residues are colored yellow. The positions of residue in green are crucial for the substrate preference of either triose-phosphate or 3-PGA. The purple residues are critical for the specificity of PEP. **(B)** The similarity and identity of the protein sequences between Chlamydomonas pPTs and Arabidopsis TPT1. The similarity and identity were calculated using https://www.bioinformatics.org/sms2/ident_sim.html. **(C)** mRNA levels encoding putative chloroplast envelope localized pPTs under LL during growth in TP medium. **(D)** Expression of pPTs over the diel cycle; data was extracted from Zones et al., 2015. Supports Fig. 1 and 2.

**Supplementary Fig. 2:**
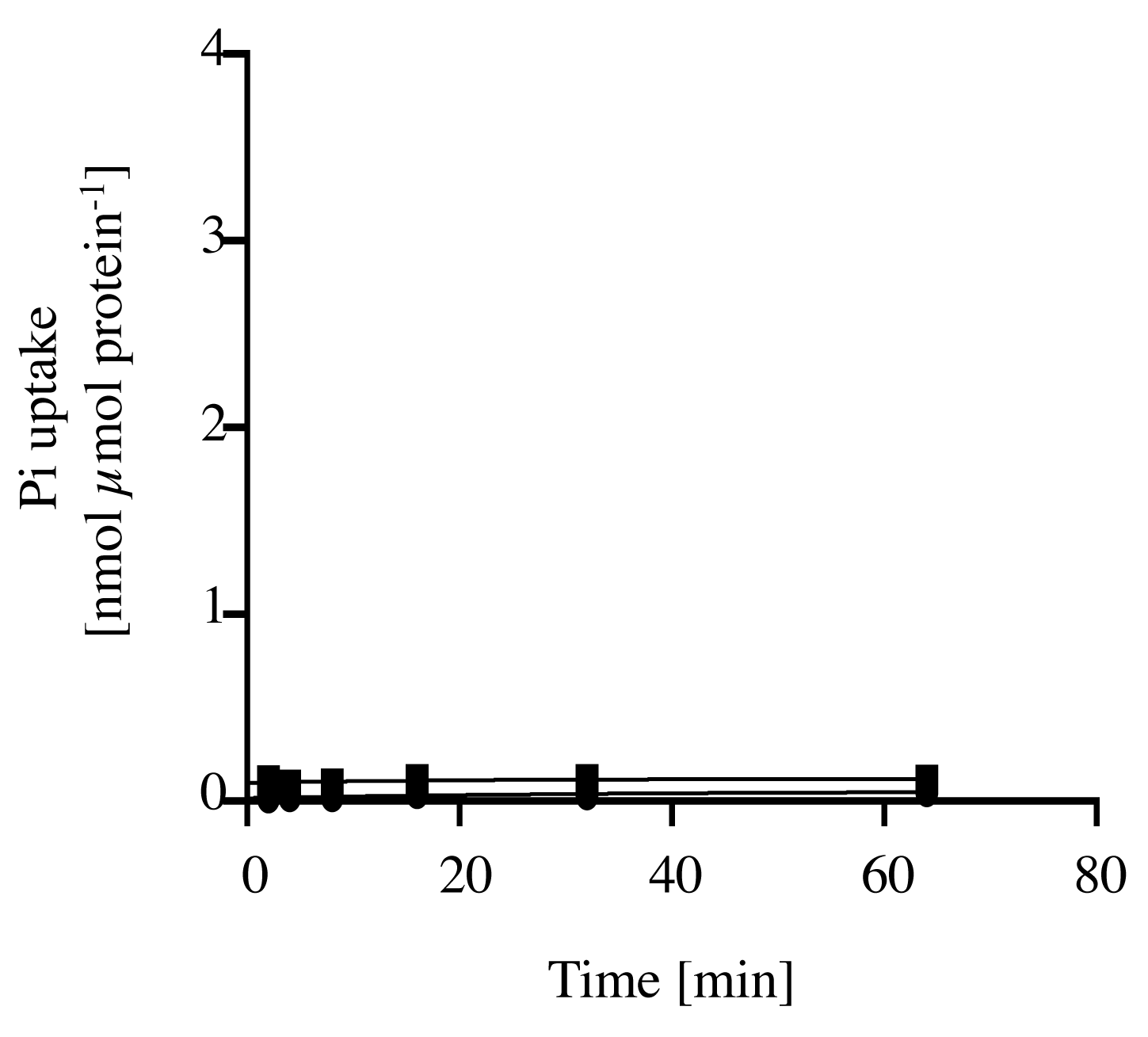
Background activity of Pi uptake with the endogenous yeast transporters reconstituted in into liposomes. Time-dependent uptake of Pi (0.25 mM) into reconstituted liposomes preloaded with 30 mM of Pi (◼) or without exchange substrate (●) prepared from yeast cells harboring the empty vector (pYES-NTa). The arithmetic mean (±SD) of three independent experiments (each with three technical replicates) was plotted against time. The observed Pi uptake rates of endogenous yeast carriers reconstituted into liposomes were negligible compared to the rates with His-CreTPT2/3 (20-fold higher, **Fig. 1C, D**), indicating that this expression system is suitable for functional analysis of recombinant phosphate transporters. Supports Fig. 1.

**Supplementary Fig. 3:**
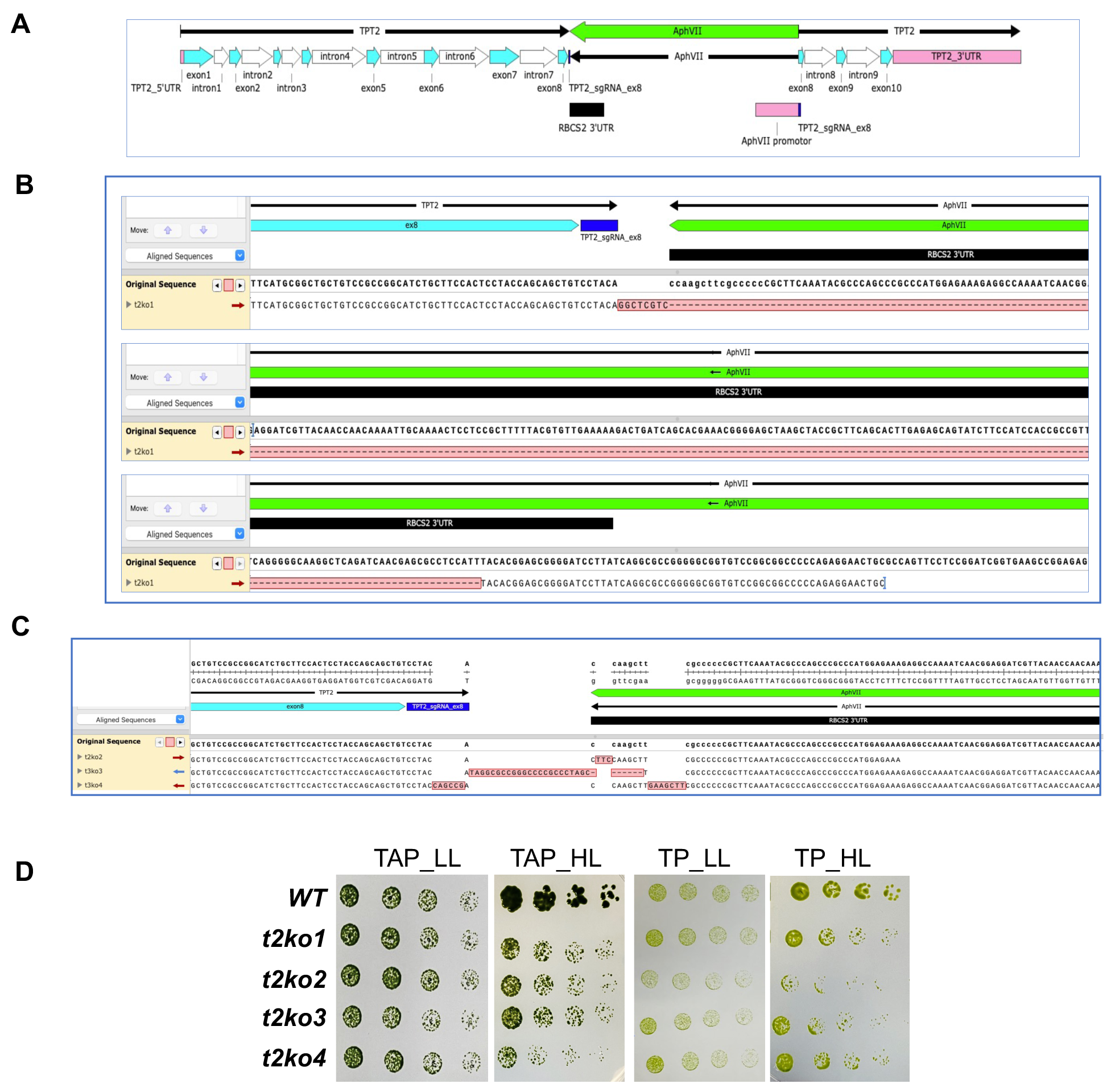
Mutations generated in the Cre*TPT2* gene by CRISPR-Cas9 mediated insertion. (**A**) Presentation of the orientation and position of *AphVII* cassette in the Cre*TPT2* gene in *t2ko1*, *t2ko2*, *t2ko3* and *t2ko4*. Sequencing of genomic DNA fragments across the site of *AphVII* cassette in *t2ko1* (**B**) and *t2ko2, t2ko3* and *t2ko4* (**C**). The mismatched nucleotides are marked in pink. Deletions are shown with dashes. (**D**) Growth of WT, *t2ko1*, *t2ko2*, *t2ko3* and *t2ko4* at various dilutions on agar plates with TP or TAP medium and incubated under LL and HL for 6 d. The dilution series used was 1, 0.5, 0.25, 0.125 μg/mL chlorophyll (left to right). Supports Fig. 2.

**Supplementary Fig. 4:**
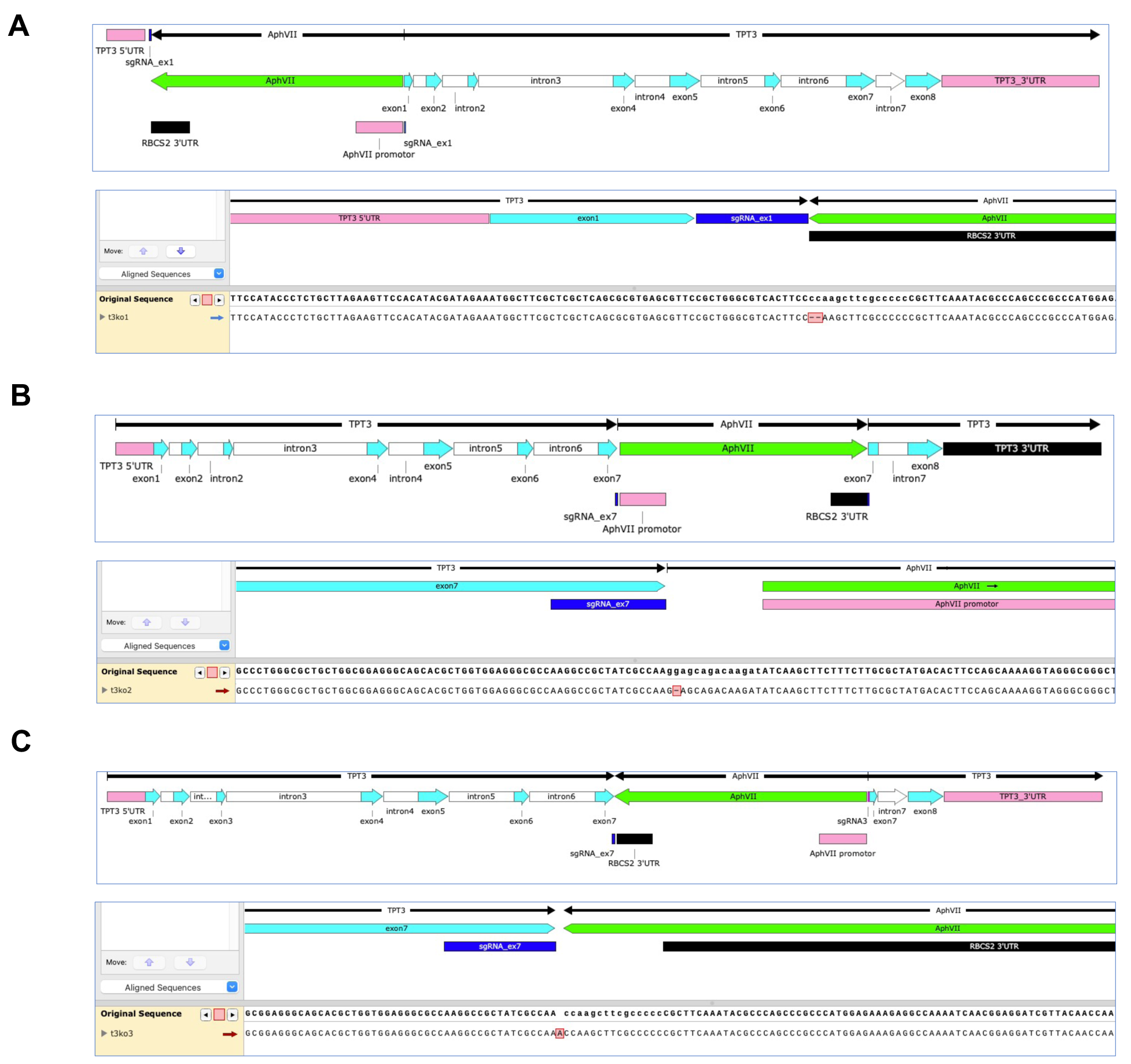
Types of mutations generated in the Cre*TPT3* gene by CRISPR-Cas9 mediated insertion with sgRNA1 in exon1 and sgRNA2 in exon7. Sequencing of genomic DNA fragments across the site of insertion of *AphVII* cassette in *t3ko1* (A) *t3ko2* (B) and *t3ko3* (C). The orientation and position of *AphVII* cassette are presented above the alignment. The mismatched nucleotides are marked in pink. Deletions are shown with dashes. Supports Fig. 2.

**Supplementary Fig. 5:**
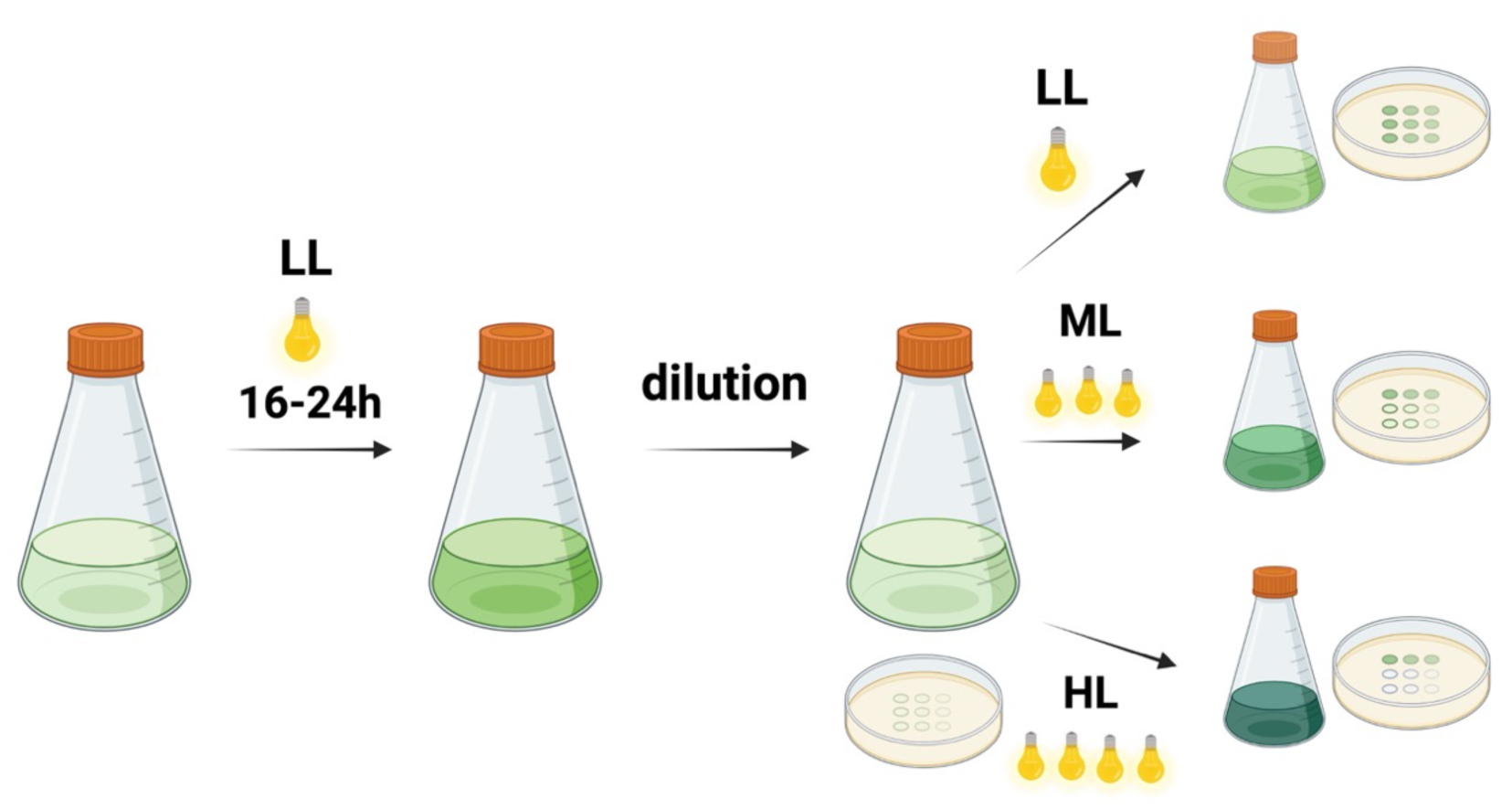
Experimental set up. All strains were grown photoautotrophically under low white light (LL, 30 µmol photons m^-2^ s^-1^) for 16-24 h. After dilution with fresh medium, the cultures from LL were then either spotted onto solid medium or to liquid medium and maintained at either LL, moderate light (ML, 250-300 μmol photon m^-2^ s^-1^) or high light (HL, 450 μmol photon m^-2^ s^-1^) for 24 or 48 h. To avoid self-shading, the culture density was adjusted to 1-1.5 µg/mL chlorophyll before exposure to ML or HL. Supports Fig. 2,3,4,5.

**Supplementary Fig. 6:**
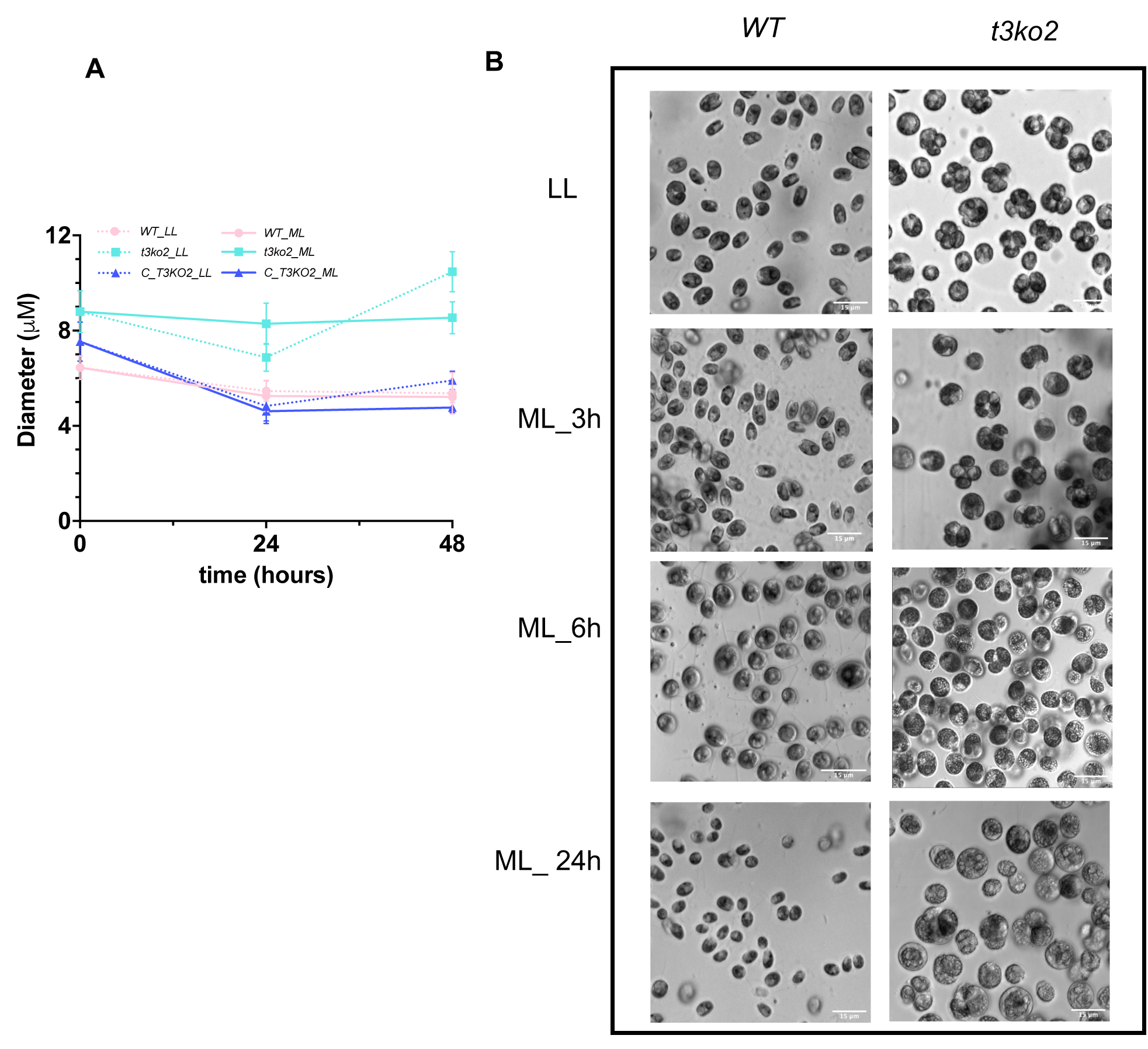
(**A**) Cell size of WT, *t3ko2*, and *C_T3KO2* in TP in both LL and ML. (**B**) WT and *t3ko2* morphology 24 h following a LL to ML transition. Supports Fig. 2.

**Supplementary Fig. 7:**
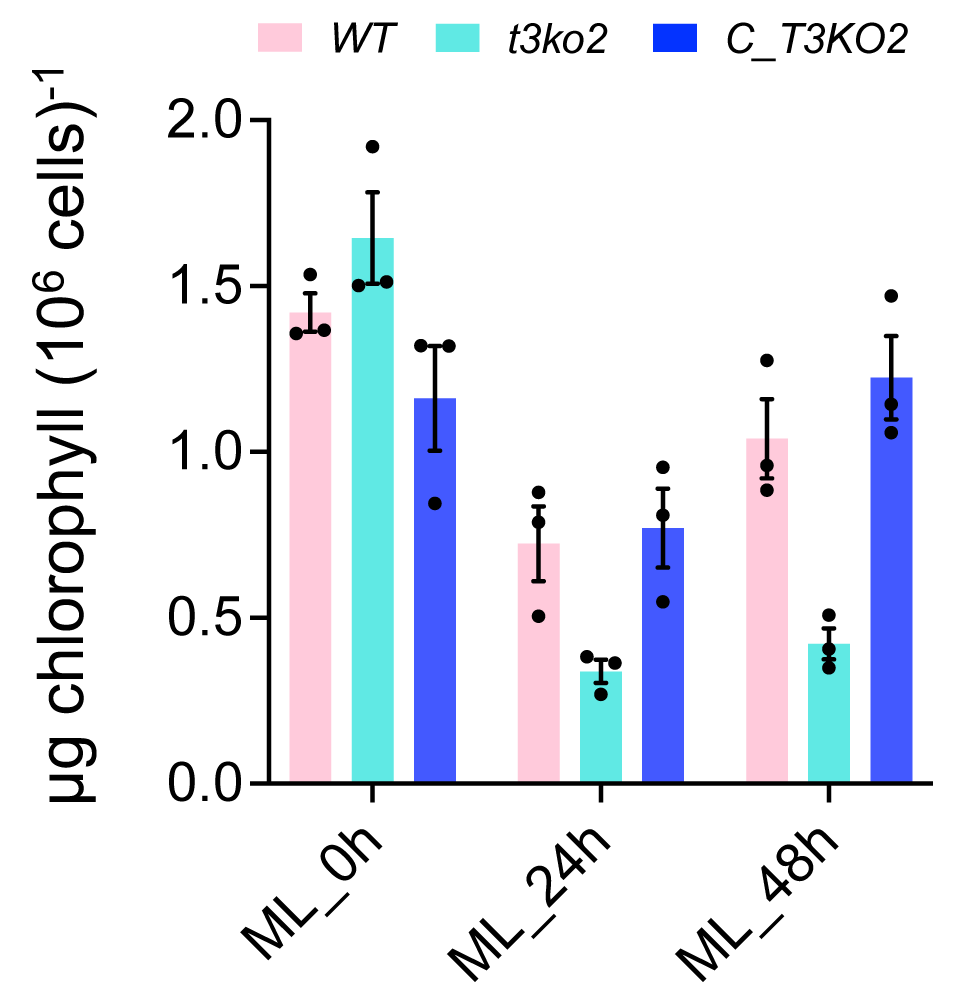
Quantification of chlorophyll contents in indicated strains after exposure to ML for 24 and 48 h. All strains were grown photoautotrophically in LL for 16-24 h. After dilution with fresh medium, the culture was transferred to ML and the chlorophyll content determined after 24 and 48 h in the ML. The initial inoculum density used for the transition was between 1-1.5 µg/mL chlorophyll in order to avoid self-shading. Supports Fig. 2.

**Supplementary Fig. 8:**
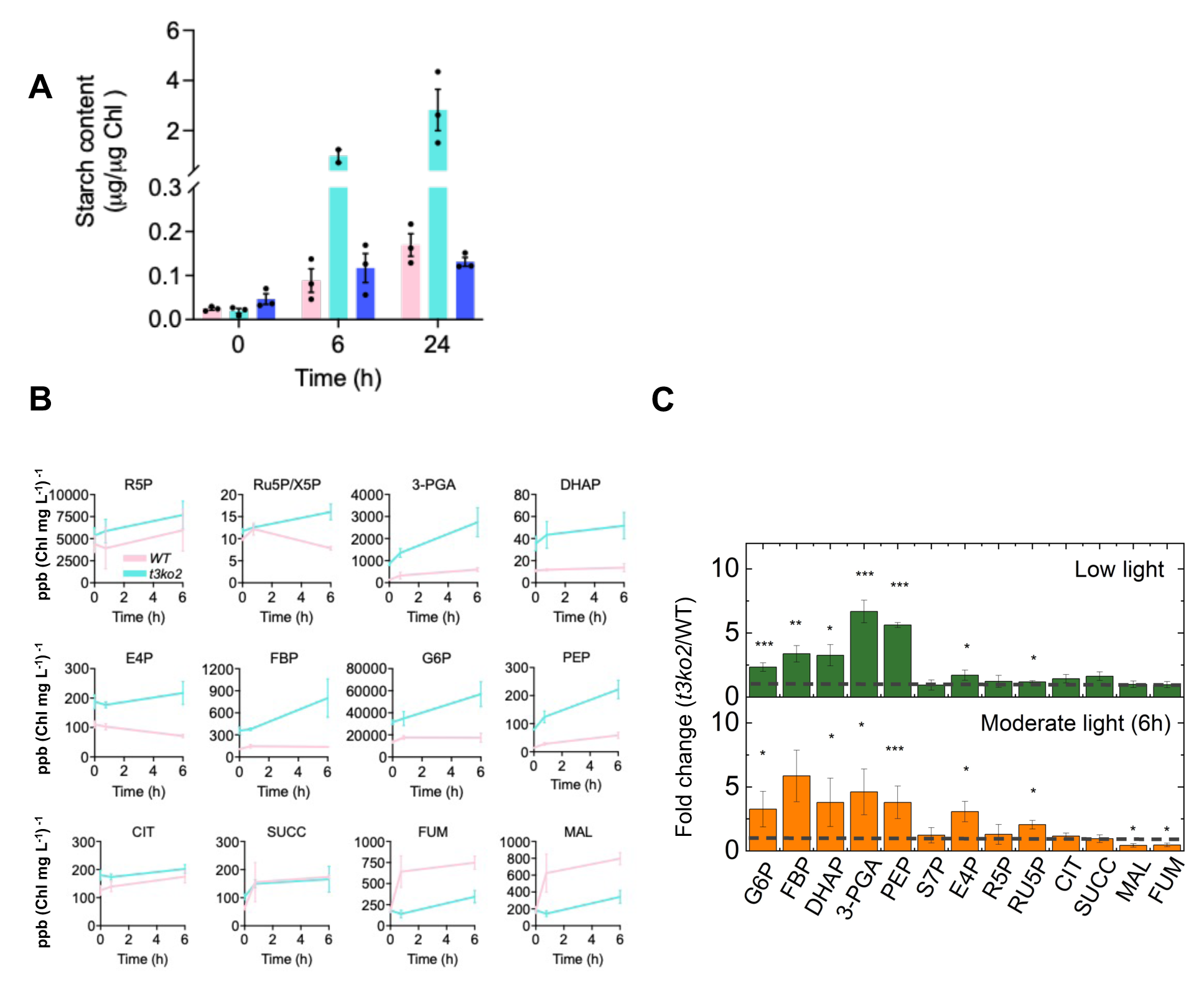
Starch and metabolites in WT and *t3ko2*. (**A)** Starch content in indicated strains following 0, 6, and 24 h in LL. (**B**) Time course, over a period of 6 h, of changes in metabolite pool sizes measured at 0 h (LL), 45 min and 6 h after shifting LL-grown cells to ML. Data was normalized to chlorophyll contents. (**C**) Schematic representation of the fold-change for the metabolites shown in (**B**), calculated by dividing the pool size in *t3ko2* by that of WT cells under the same conditions. Fold change of 1 (no change) is shown by a dashed line. Each data point is the mean and standard error of three biological replicates. ppb: parts per billion. Asterisk indicates statistically significant differences relative to WT (* P<0.05, ** P<0.01, *** P<0.001). Abbreviations: G6P, glucose-6-phosphate; FBP, fructose bisphosphate; DHAP, dihydroxyacetone phosphate; 3-PGA, 3-phosphoglycerate; PEP, phosphoenolpyruvate; CIT, citrate; SUCC, succinate; FUM, fumarate; MAL, malate; S7P, sedoheptulose-7-phosphate; E4P, Erythrose 4-phosphate; R5P, Ribose 5-phosphate; RU5P/X5P, ribulose 5-phosphate/xylulose-5-phosphate. Supports Fig. 3.

**Supplementary Fig. 9:**
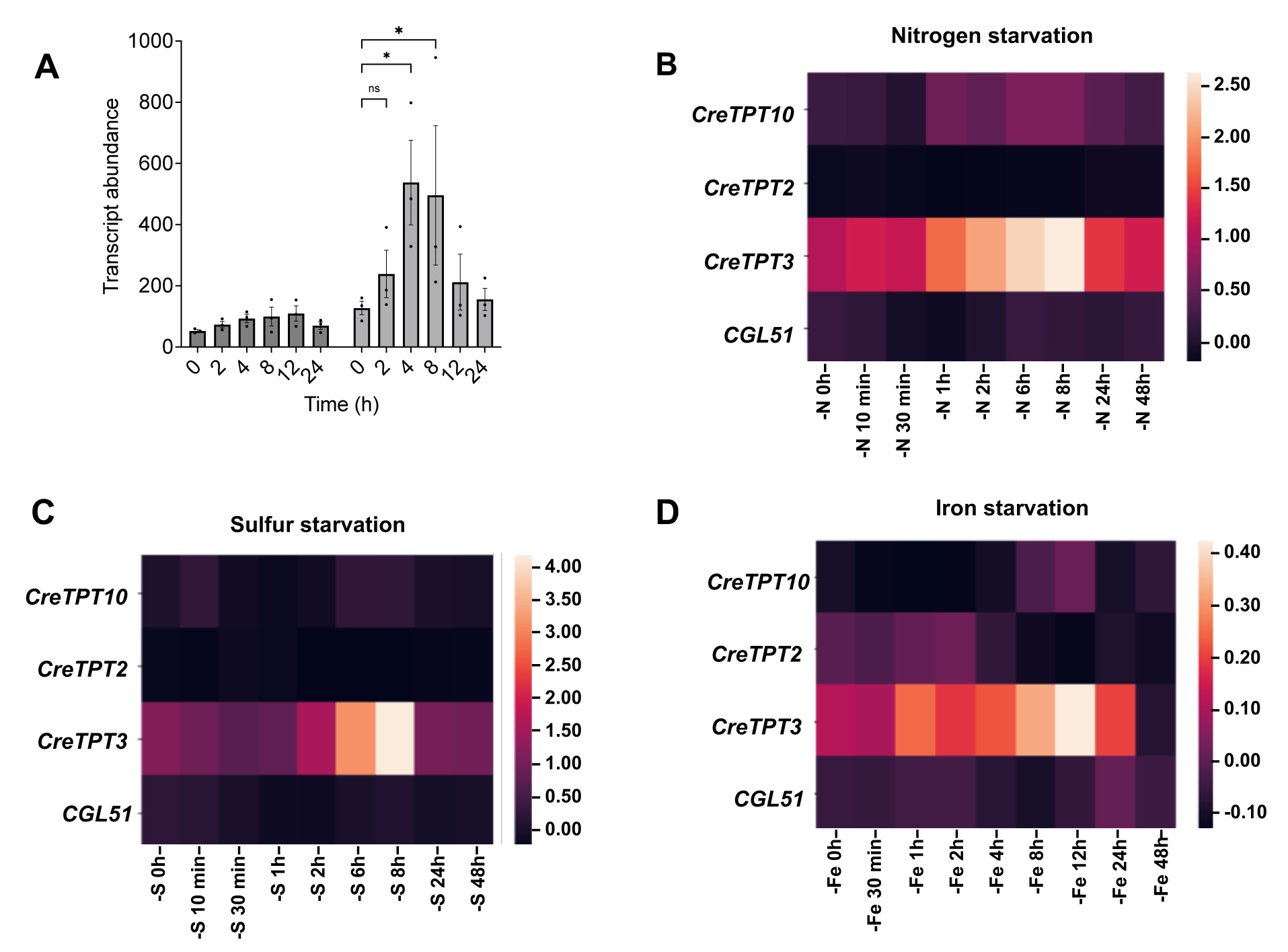
transcript changes of Cre*TPT2* and Cre*TPT3* with various conditions. **(A)** Expressions of Cre*TPT2* and Cre*TPT3* following exposure of LL grown cells to HL. Expression of the putative pPTs under nitrogen starvation **(B)**, sulfur starvation **(C)**, and iron starvation **(D)** from (Ngan et al., 2015; González-Ballester et al., 2010; Urzica et al., 2013). Supports Fig. 5,6,7.

**Supplementary Fig. 10:**
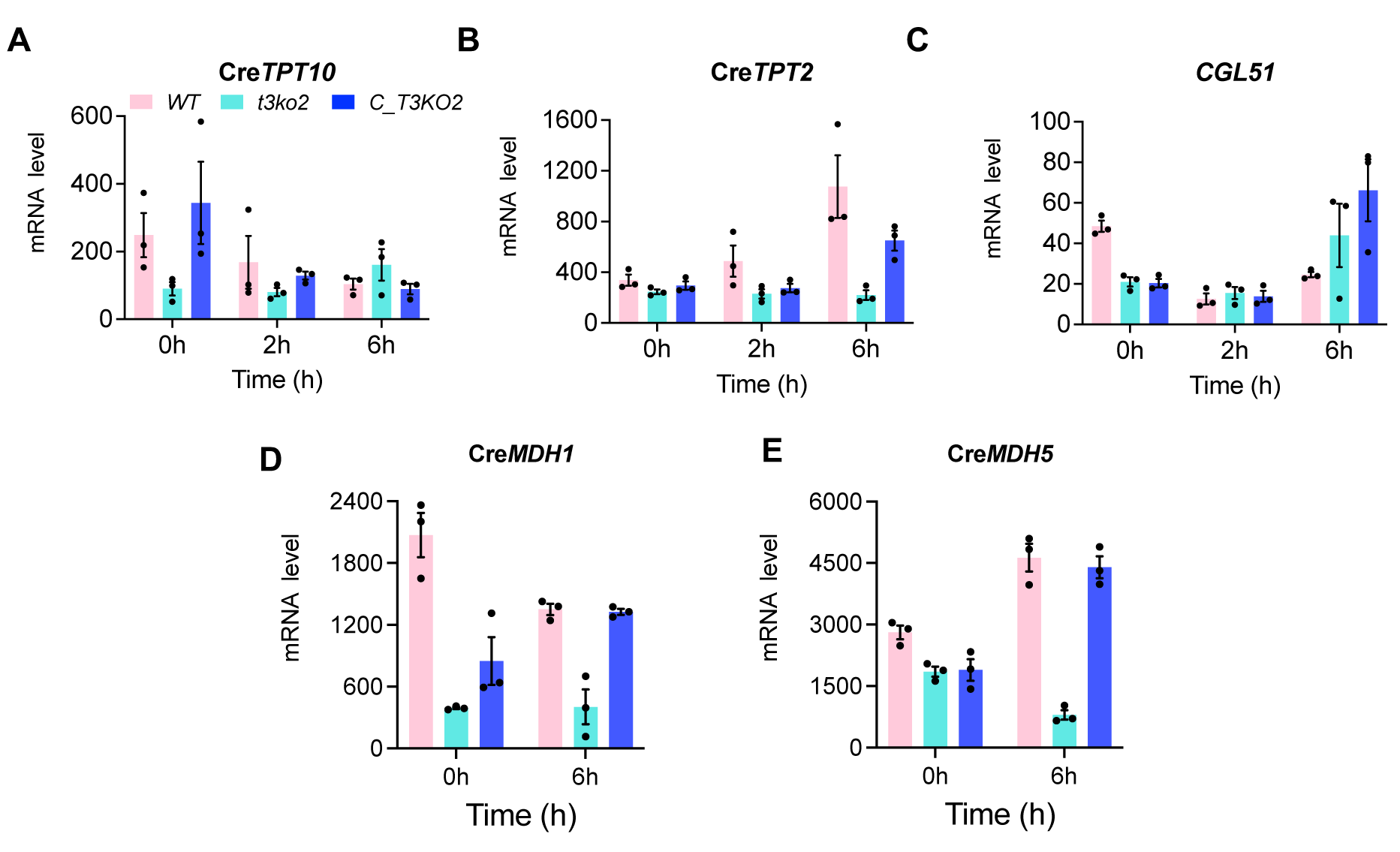
mRNA levels of Cre*TPT10* (**A**), Cre*TPT2* (**B**) and *CGL51* **(C**) in the indicated strains 0, 2, 6 h after the cells were shifted from LL to ML. (**D, E**) mRNA levels of plastidial malate dehydrogenases (*MDH*) in the indicated strains 0, 6 h after the cells were shifted from LL to ML. Supports Fig. 7.

**Supplementary Fig 11:**
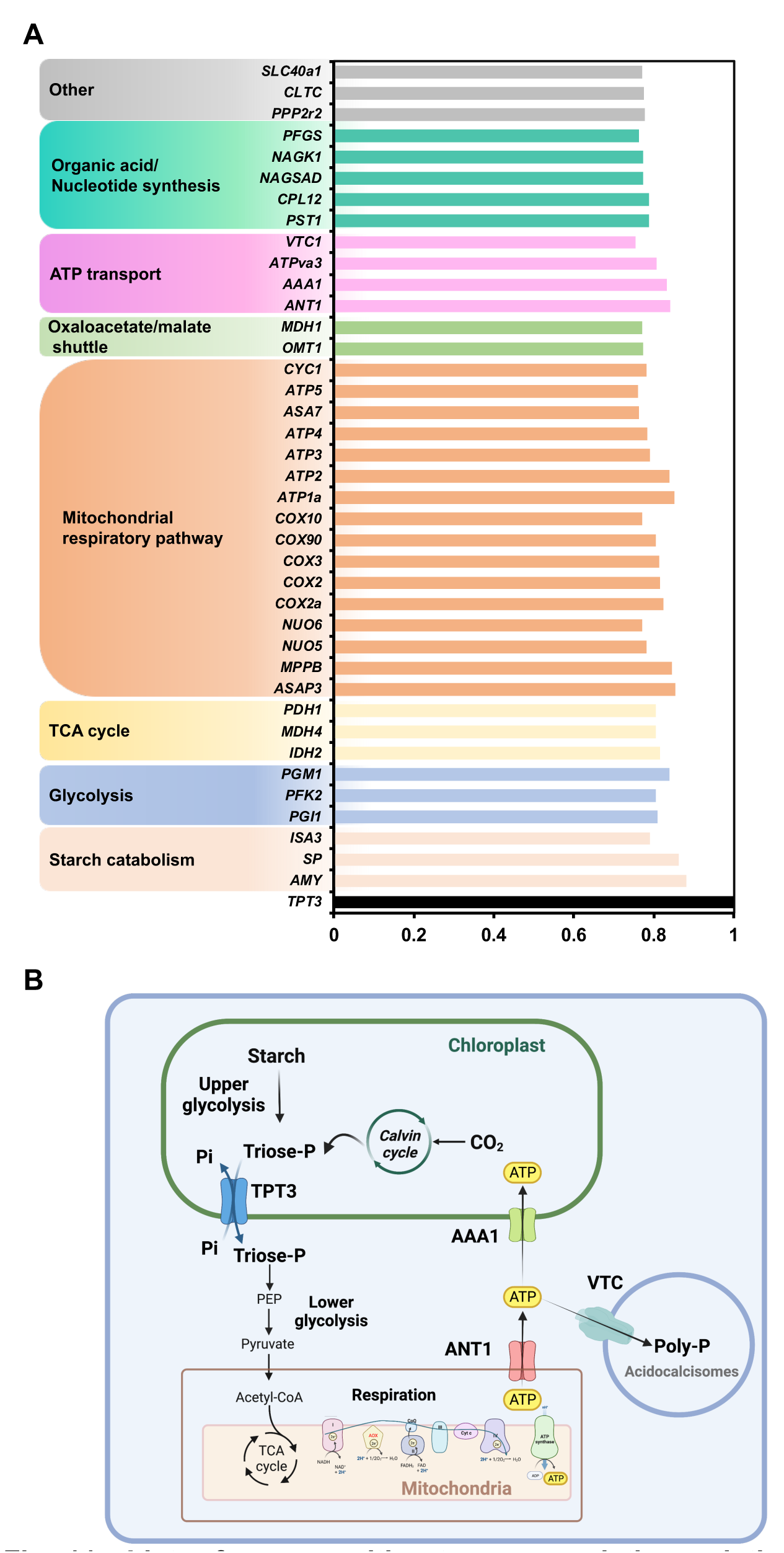
List of genes with strong correlation relative to Cre*TPT3*. (**A**) Correlation coefficients greater than 0.75 were listed. (**B**) Construction of Cre*TPT3* co-expression networks. Supports Fig. 7 and 8.

## Supplementary Tables

**Supplementary Table 1:** Members of the triose phosphate transporter family in *Chlamydomonas reinhardtii* and their predicted subcellular localization based on TargetP-2.0. cTP, chloroplast transit peptide; CS position, cleavage site of transit peptides; N: genes without names in the genome. Cre IDs are from v6.1 genome.

| Cre ID | Name from V6.1 /<br>(name used in<br>this study) | Previous name | Predicti<br>on | CS Position |
| --- | --- | --- | --- | --- |
| Cre08.g379350_4532 | TPT10 / (TPT10) | TPT1 | cTP | CS pos: 61-62. TCL-AV. Pr: 0.3994 |
| Cre06.g263850_4532 | N / (TPT2) | PPT2; TPT2; TPT20 | cTP | CS pos: 70-71. VCQ-AA. Pr: 0.5253 |
| Cre01.g045550_4532 | TPT3 / (TPT3) | TPT3 | cTP | CS pos: 60-61. VTK-AS. Pr: 0.4178 |
| Cre02.g106200_4532 | N | TPT4; TPT5 | OTHER |  |
| Cre02.g112900_4532 | N | TPT5; TPT6 | OTHER |  |
| Cre02.g144300_4532 | N | TPT6; TPT7 | OTHER |  |
| Cre03.g162000_4532 | N | TPT7; TPT8 | OTHER |  |
| Cre03.g184850_4532 | N | TPT8; TPT9 | OTHER |  |
| Cre04.g227450_4532 | N | TPT9 | OTHER |  |
| Cre07.g330850_4532 | N | TPT10; TPT11 | OTHER |  |
| Cre08.g363600_4532 | N | TPT11; TPT12 | OTHER |  |
| Cre09.g408400_4532 | N | TPT12; TPT13 | OTHER |  |
| Cre09.g413700_4532 | N | TPT13; TPT14 | OTHER |  |
| Cre08.g382350_4532 | EZY14 | TPT14; TPT16; EZY14 | OTHER |  |
| Cre09.g415900_4532 | MOT20 | TPT15 | OTHER |  |
| Cre10.g452750_4532 | N | TPT16; TPT18 | OTHER |  |
| Cre11.g479950_4532 | CGL7 | TPT17 | OTHER |  |
| Cre12.g490050_4532 | N | TPT18; TPT19; GMT1 | OTHER |  |
| Cre12.g490100_4532 | TPT2 | TPT19; GMT2 | OTHER |  |
| Cre12.g501000_4532 | N | TPT20; TPT22; PPT1 | OTHER |  |
| Cre15.g641266_4532 | N | TPT22; TPT23 | OTHER |  |
| Cre15.g642950_4532 | N | TPT23; TPT24 | OTHER |  |
| Cre15.g643385_4532 | N | TPT24; TPT26 | OTHER |  |
| Cre16.g663800_4532 | CGL51 / (CGL51) | TPT25 | cTP | CS pos: 70-71. IVA-SS. Pr: 0.4817 |
| Cre16.g666250_4532 | N | TPT26; TPT27 | OTHER |  |
| Cre17.g702700_4532 | N | TPT27; TPT28 | OTHER |  |
| Cre17.g703250_4532 | SLC35D | TPT28; TPT29 | OTHER |  |
| Cre17.g710850_4532 | N | TPT29; TPT4 | OTHER |  |
| Cre11.g467754_4532 | N | UAA6 | OTHER |  |
| Cre14.g622700_4532 | N | N | OTHER |  |
| Cre18.g748947_4532 | N | N | OTHER |  |
| Cre09.g408428_4532 | N | N | OTHER |  |

**Supplementary Table 2:**
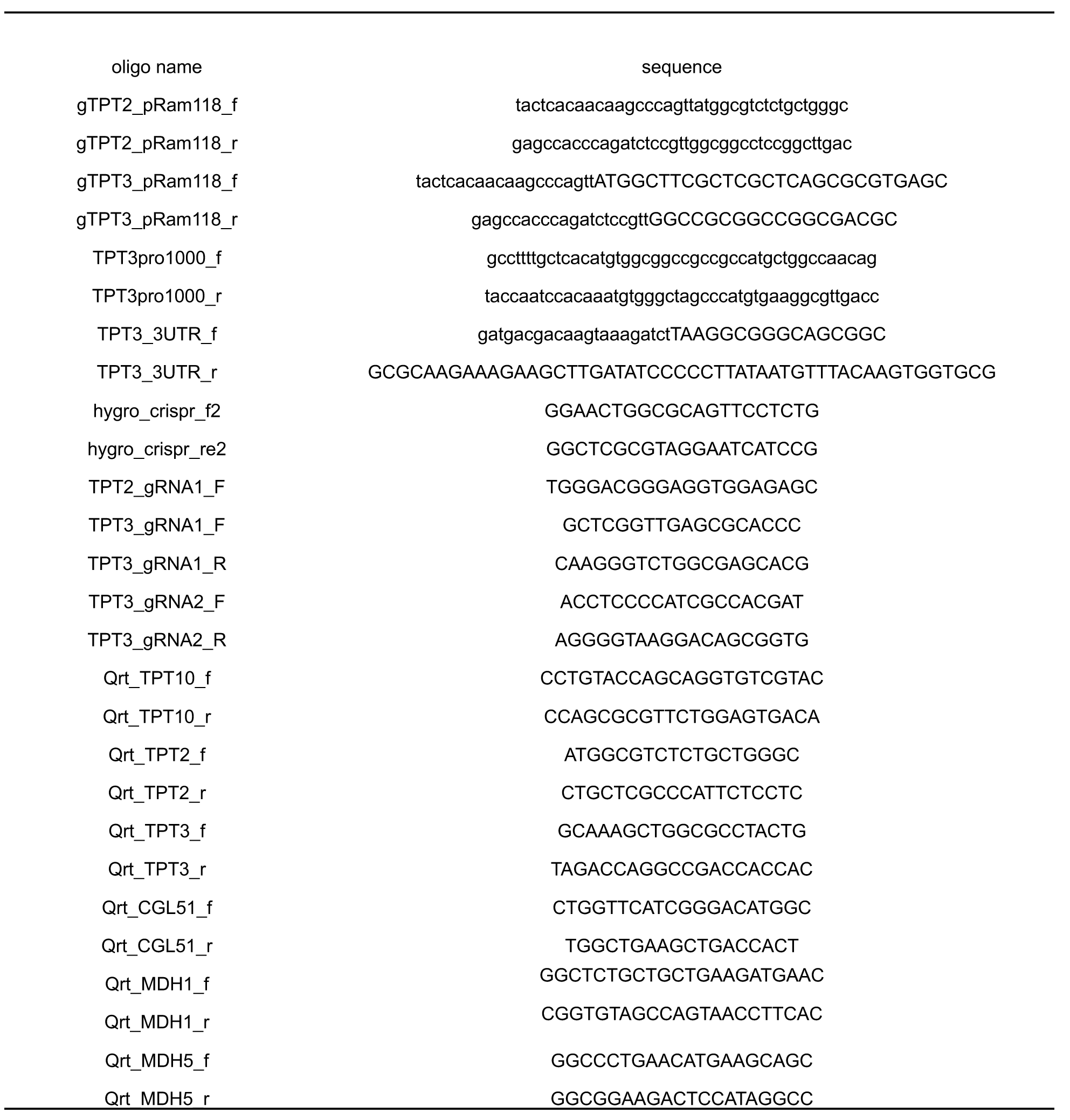
Oligonucleotide list.

**Supplementary Table 3:**
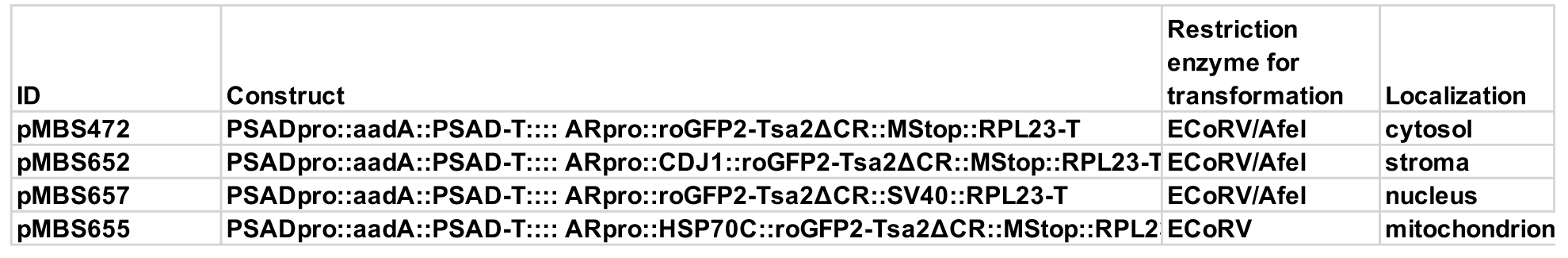
Plasmids of H_2_O_2_ roGFP2-Tsa2ΔCR sensors (Niemeyer et al., 2021).

**Supplementary Table 4:** List of genes with strong Pearson correlation to *CreTPT3*.1.

| **Name** | **Cre number** | **Gene function** | **Correlation coefficient** | **Localization** |
| --- | --- | --- | --- | --- |
| *TPT3* | Cre01.g045550 | Triose phosphate/phosphate translocator 2 | 1 | chloroplast |
| *AMY* | Cre08.g362450 | Alpha-amylase | 0.881792677 | chloroplast |
| *SP* | Cre07.g336950 | Starch phosphorylase | 0.861504451 | chloroplast |
| *ISA3* | Cre03.g207713 | Isoamylase, starch debranching enzyme | 0.790501386 | chloroplast |
| *PGI1* | Cre03.g175400 | Phosphoglucose isomerase | 0.808137223 | chloroplast |
| *PFK2* | Cre12.g553250 | Phosphofructokinase | 0.805294876 | chloroplast |
| *PGM1* | Cre06.g272050 | Phosphoglycerate mutase | 0.838232423 | other |
| *IDH2* | Cre02.g143250 | Isocitrate dehydrogenase, NAD-dependent | 0.81622058 | other |
| *MDH4* | Cre12.g483950 | Malate dehydrogenase 4 | 0.80363604 | mitochondrion |
| *PDH1* | Cre16.g677026 | Pyruvate dehydrogenase E1 beta subunit | 0.805186998 | mitochondrion |
| *ASAP3* | Cre07.g338050 | Mitochondrial F1F0 ATP synthase associated protein 3 | 0.853067434 | mitochondrion |
| *MPPB* | Cre12.g523850 | Mitochondrial processing peptidase beta subunit and Complex III Core I subunit | 0.844453019 | mitochondrion |
| *NUO5* | Cre10.g450400 | NADH dehydrogenase (ubiquinone) flavoprotein 2, mitochondrial (respiratory Complex I) | 0.781322987 | mitochondrion |
| *NUO6* | Cre10.g422600 | NADH:ubiquinone oxidoreductase 51 kDa subunit, mitochondrial, respiratory Complex I | 0.770479958 | mitochondrion |
| *COX2a* | Cre03.g154350 | Mitochondrial cytochrome c oxidase subunit II | 0.824639841 | mitochondrion |
| *COX2* | Cre01.g049500 | Mitochondrial cytochrome c oxidase subunit II, protein IIb of split subunit | 0.814345856 | mitochondrion |
| *COX3* | Cre04.g221700 | Mitochondrial cytochrome c oxidase subunit III | 0.813942712 | mitochondrion |
| *COX90* | Cre16.g691850 | Cytochrome c oxidase subunit Cox90, mitochondrial | 0.803959451 | mitochondrion |
| *COX10* | Cre12.g516350 | Mitochondrial cytochrome c oxidase assembly protein | 0.770332065 | mitochondrion |
| *ATP1a* | Cre02.g116750 | Mitochondrial F1F0 ATP synthase, alpha subunit | 0.850554844 | mitochondrion |
| *ATP2* | Cre17.g698000 | Mitochondrial F1F0 ATP synthase, beta subunit | 0.839116637 | mitochondrion |
| *ATP3* | Cre15.g635850 | Mitochondrial F1F0 ATP synthase, gamma subunit | 0.790390859 | mitochondrion |
| *ATP4* | Cre11.g467707 | Mitochondrial F1F0 ATP synthase, delta subunit | 0.782621502 | mitochondrion |
| *ASA7* | Cre09.g416150 | Mitochondrial F1F0 ATP synthase associated protein 7 | 0.762529504 | mitochondrion |
| *ATP5* | Cre16.g680000 | Mitochondrial ATP synthase subunit 5 | 0.75995889 | mitochondrion |
| *CYC1* | Cre15.g638500 | Ubiquinol:cytochrome c oxidoreductase cytochrome c1, mitochondrial | 0.78087886 | mitochondrion |
| *OMT1* | Cre17.g713350 | Oxoglutarate:malate antiporter1 | 0.7724107 | chloroplast |
| *MDH1* | Cre03.g194850 | NAD-dependent malate dehydrogenase | 0.770420148 | chloroplast |
| *ANT1* | Cre09.g386650 | ADP/ATP carrier protein, mitochondrial | 0.841663755 | mitochondrion |
| *AAA1* | Cre08.g358526 | Plastidic ADP/ATP translocase | 0.831658774 | chloroplast |
| *ATPva3* | Cre04.g220350 | Vacuolar ATP synthase subunit A | 0.806662753 | other |
| *VTC1* | Cre12.g510250 | Vacuolar Transport Chaperone-like protein | 0.753558866 | other |
| *PST1* | Cre07.g331550 | Phosphoserine aminotransferase | 0.787494263 | chloroplast |
| *CPL12* | Cre10.g466500 | Glyoxalase | 0.787085275 | chloroplast |
| *NAGSAD* | Cre03.g146187 | N-acetyl-gamma-glutamyl-phosphate reductase / NAGSA dehydrogenase/oxidoreductase | 0.772201917 | chloroplast |
| *NAGK1* | Cre01.g015000 | Acetylglutamate kinase/L-arginine biosynthesis II (acetyl cycle) | 0.771816595 | chloroplast |
| *PFGS* | Cre08.g364800 | Phosphoribosylformylglycinamide synthase /superpathway of purine nucleotides | 0.761886415 | chloroplast |
| *PPP2r2* | Cre01.g055420 | serine/threonine-protein phosphatase 2A regulatory subunit B | 0.776264136 | chloroplast |
| *CLTC* | Cre02.g101400 | clathrin heavy chain | 0.775380317 | other |
| *SLC40a1* | Cre06.g251000 | solute carrier family 40 | 0.771233669 | secretory pathway |

